# Gene loss and cis-regulatory novelty shaped core histone gene evolution in the apiculate yeast *Hanseniaspora uvarum*

**DOI:** 10.1101/2023.08.28.551515

**Authors:** Max A. B. Haase, Jacob L. Steenwyk, Jef D. Boeke

## Abstract

Across eukaryotic species, core histone genes display a remarkable diversity of cis-regulatory mechanisms despite their protein sequence conservation. However, the dynamics and significance of this regulatory turnover are not well understood. Here we describe the evolutionary history of core histone gene regulation across 400 million years in Saccharomycotina yeasts, revealing diverse lineage-specific solutions to core histone expression. We further characterize the emergence of a novel regulatory mode in the *Hanseniaspora* genus, which coincided with the loss of one copy of its paralogous core histones genes from its fast-evolving lineage. By analyzing the growth dynamics using live cell imaging of genetically modified *Hanseniaspora uvarum*, we observed a regulatory decoupling of core histone synthesis from DNA synthesis and propose that this may be adaptive for its rapid cell cycle progression. Overall, our findings imply that the frequent turnover of core histone cis-regulatory mechanisms likely provides distinct adaptive solutions for specific life histories.

## Introduction

Chromatin structure and function are critical for essential processes, including DNA replication, chromosome division, DNA damage repair, and transcription (Wei *et al*. 1999; MacAlpine and Almouzni 2013; Venkatesh and Workman 2015; Hauer *et al*. 2017; Haase *et al*. 2023a; b; Lazar-Stefanita *et al*. 2023). The basic unit, the core nucleosome particle, is an octameric protein complex made up of an H3-H4 tetramer flanked by two separate H2A-H2B dimers that together wrap ∼146 bp of DNA (Luger *et al*. 1997). Core histone genes are often present at multiple copies encoded as gene clusters in eukaryotic species. In the yeast *Saccharomyces cerevisiae,* each core histone is encoded by two paralogous genes that are arranged in the genome as divergently transcribed clusters, either encoding H2A-H2B (*HTA1B1* and *HTA2B2* loci) or H3-H4 (*HHF1T1* and *HHF2T2* loci; Eriksson *et al*. 2012). Reflecting their fundamental importance, the nucleosome structure and the primary amino acid sequence of histones are highly conserved across eukaryotes (Malik and Henikoff 2003); the yeast paralogous histone proteins are identical (H3/4 and H2B) or near identical (H2A). Tight control of the regulation of core histones throughout the cell cycle ensures the proper function of DNA-templated processes (Eriksson *et al*. 2012). Proper stoichiometry between nucleosomes and total DNA content is partly controlled through replication-coupled synthesis of histones (Robbins and Borun 1967), evidenced by experimental inhibition of DNA synthesis leading to rapid repression of histone synthesis (Osley 1991; Rattray and Müller 2012; Bhagwat *et al*. 2021) Moreover, misexpression of histones outside of S-phase characteristically leads to cellular toxicity and growth arrest (Kurat *et al*. 2014).

Despite the evolutionarily conserved nature of cycle-dependent expression and function of histones (Jensen *et al*. 2006), the cis-regulatory mechanisms used to achieve precise control of core histone expression are diverse across eukaryotes (Mariño-Ramírez *et al*. 2006). In the budding yeast *Saccharomyces cerevisiae*, specific S-phase expression is driven by both positive and negative regulatory mechanisms (Eriksson *et al*. 2012; Kurat *et al*. 2014). The primary control is mediated by the transcription factor Spt10, a putative acetyltransferase that positively induces transcription through the histone upstream activating sequence (UAS; Dollard *et al*. 1994; Eriksson *et al*. 2005). Prior work has shown that this Spt10-mode is deeply conserved across yeast phylogeny (Mariño-Ramírez *et al*. 2006); however, the specific origins and emergence of this regulatory mode still need clarification. In addition, negative regulatory feedback is achieved through various mechanisms (Eriksson *et al*. 2012), including various histone chaperones (HIR complex, Atf1, and Rtt106), RSC chromatin remodelers, and by less well-conserved DNA-protein interactions (i.e., NEG region; Eriksson *et al*. 2012). Among budding yeasts, genes encoding histones typically lack introns (Yun and Nishida 2011), suggesting splicing does not play a prominent role in post-transcriptional regulation. Despite these advances, the specific origins and evolutionary outcomes of histone regulatory modes in budding yeasts remain largely unknown.

Mechanisms of core histone regulation are particularly enigmatic in the genus *Hanseniaspora*, a group of bipolar budding yeasts belonging to the order Saccharomycodales (Groenewald *et al*. 2023). The evolutionary history of *Hanseniaspora* spp. is marked by a burst of rapid evolution and the loss of numerous conserved genes, including those associated with cell cycle processes and genes involved in DNA repair (Steenwyk *et al*. 2019). As a result, some *Hanseniaspora* spp. have genome sizes of around 8-9 megabases and encode approximately 4,000 genes (Steenwyk *et al*. 2019). In contrast, *S. cerevisiae* has a genome size of roughly 12 megabases and encodes approximately 6,000 genes (Goffeau *et al*. 1996). The degree of rapid evolution gene loss is more pronounced in one *Hanseniaspora* lineage compared to the other and are thus termed the faster-evolving lineage (FEL) and slower-evolving lineage (SEL), respectively. However, the dearth of established tools for genetic manipulation of *Hanseniaspora* spp. has stymied our understanding of its cell biology and genetics (Schwarz *et al*. 2022; Heinisch *et al*. 2023), including how gene loss has (re)shaped cell cycle processes. In contrast, much more is known about *Hanseniaspora* ecology. *Hanseniaspora* spp. are abundantly present on various fruits (e.g., grapes) and associate with various insects (such as *Drosophila* spp.) attracted to volatile aromatic compounds produced by *Hanseniaspora spp.* (Hamby *et al*. 2012; Becher *et al*. 2018; Saubin *et al*. 2020)*. Hanseniaspora* has also gained interest in biotechnology applications, such as expanding the sensorial complexities of fermented products (Steensels and Verstrepen 2014) and as natural biocontrol agents (Rueda-Mejia *et al*. 2022). Among *Hanseniaspora* spp., *Hanseniaspora uvarum*, a species in the FEL, has become a focus for research and biotechnological development (Badura *et al*. 2021, 2023; Heinisch *et al*. 2023; Van Wyk *et al*. 2023).

Here, we explore the evolution of core histone genes and their cis-regulatory evolution in a panel of yeast that span the diversity of the Saccharomycotina subphylum and conduct an in-depth computational and molecular investigation of *Hanseniaspora*. Using a histone replacement assay, we find evidence suggesting that *H*. *uvarum*’s H2A-H2B dimer and, surprisingly, its histone promoters are incompatible in *S*. *cerevisiae*. Examination of the cis-regulatory changes underlying the histone promoter incompatibility revealed that the ancestral Spt10-mode was replaced with a derived Mcm1-mode of regulation in *H*. *uvarum* and other FEL species. Moreover, we show that the function and regulatory network of *H*. *uvarum*’s Mcm1 is conserved with *S*. *cerevisiae*, suggesting that the histones were rewired into a novel regulatory paradigm in *H*. *uvarum*. Characterizing cell cycle dynamics and the timing of histone synthesis in single cells of *H*. *uvarum,* we uncovered a rapid division with a doubling time of ∼60 minutes, and surprisingly, we found that histone and DNA synthesis status were decoupled, unlike the case in *S. cerevisiae*. These findings uncovered unexpected novelty in a hitherto conserved and fundamental cellular process. More broadly, this work lays the foundation for future genetic investigations into the highly divergent and rapidly dividing genus of *Hanseniaspora*.

## Results

### Paralogous core histone gene loss and divergence in *Hanseniaspora* FEL

Members of the bipolar budding yeast genus *Hanseniaspora* have dramatically reduced genomes lacking many essential cell-cycle and repair regulators (Steenwyk *et al*. 2019). This is reflected between two lineages– the slower-evolving and faster-evolving lineages (SEL and FEL), with FELs, including *H. uvarum* studied here, showing the most dramatic levels of gene loss (Steenwyk *et al*. 2019). However, an analysis of *Hanseniaspora* core histones has yet to be reported. To investigate the evolution of core histones in *Hanseniaspora*, we first set out to detail the copy number evolution of histone genes. In the out-group species and *Hanseniaspora* SEL species, we observed the presence of the canonical paralogous histone gene clusters, as in *S. cerevisiae* (Figure 1A, Figure S1). However, for species of the *Hanseniaspora* FEL, we observed only a single copy of each histone gene cluster (Figure 1A). Synteny analysis, suggests the that the histone clusters *HTA2B2* and *HHF1T1* have been lost in the *Hanseniaspora* FEL ancestor (Figure S1B–C). Intriguingly, we also observed a convergent partial loss event in one species of the *Hanseniaspora* SEL, *H. gamundiae*, which lost paralog cluster *HHT1F1,* suggesting it may represent an independent intermediate state or an artifact of incomplete genome sequencing and assembly (Figure 1A, Figure S1A).

**Figure 1.**
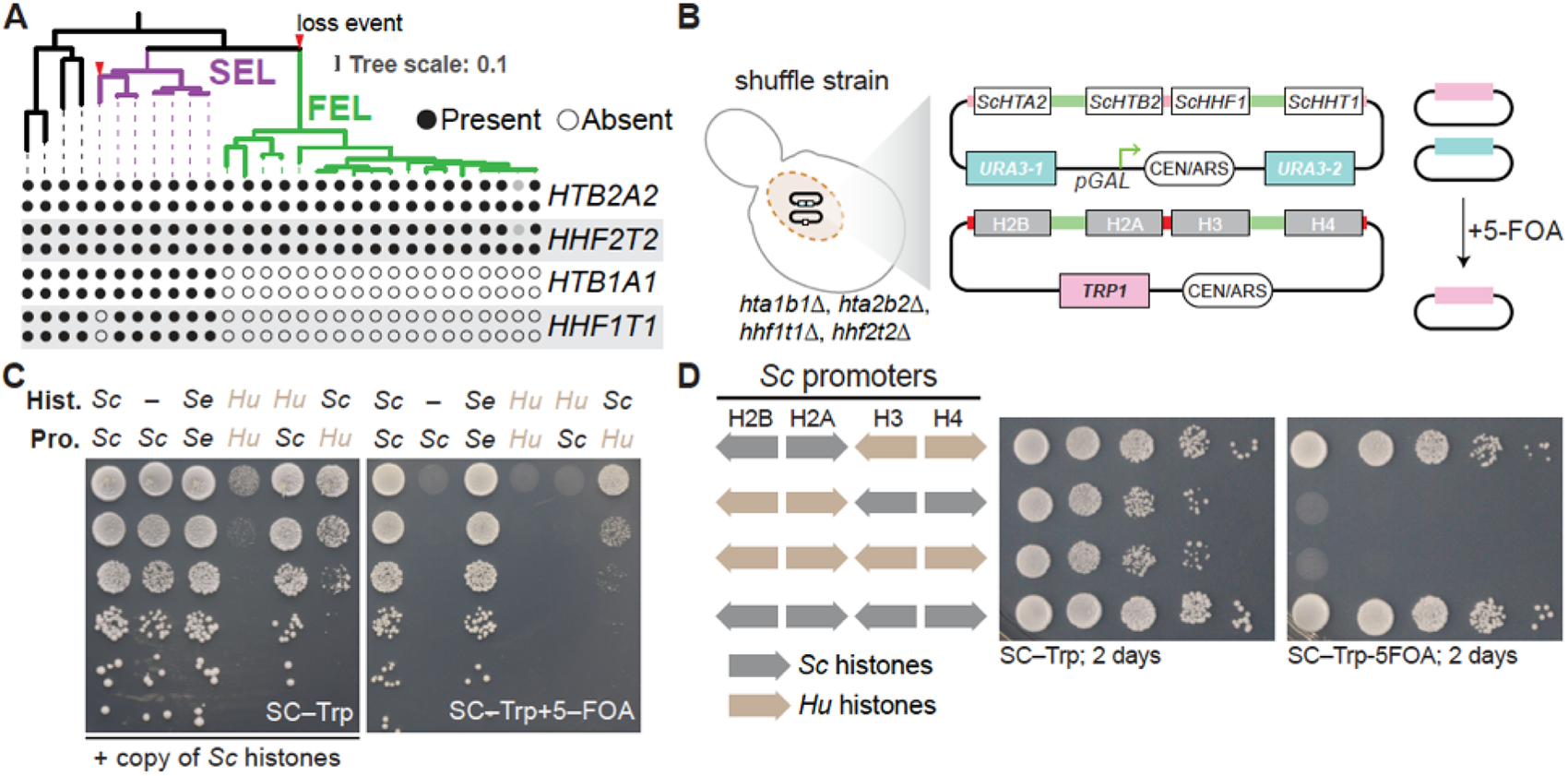
Paralogous gene loss and divergence of core histones in *Hanseniapsora* FEL. **A**. Phylogeny of the *Hanseniaspora* and four outgroup species from Steenwyk et al. (2019), showing the presence and absence of the core histones gene clusters. Purple, the slow-evolving lineage; Green, the fast-evolving lineage. Outgroup lineages from left to right: *S*. *cerevisiae*, *K*. *lactis*, *Cyberlindnera jadinii*, *Wickerhamomyces anomalus*. The full phylogeny, complete with species and strain names, can be found in Supplementary Figure 1A. **B**. Overview of dual-plasmid histone shuffle assay in *S*. *cerevisiae*. Details of plasmid shuffling can be found in the methods and also in Haase *et al*. 2019, 2023a. **C**. Shuffle assay of *H*. *uvarum*’s histones and histone gene promoters in *S. cerevisiae*. Left is the growth assay maintaining the selection for both plasmids, and right is the counterselection (5–FOA) of the native *S*. *cerevisiae* histone plasmid. Abbreviations are used for displaying which host species each histone promoter and genes were sourced from Sc – *S*. *cerevisiae*; Se – *S. eubayanus*; Hu – *H*. *uvarum*. **D**. Shuffle assay of *H*. *uvarum*’s H2A-H2B and H3-H4 dimers in *S. cerevisiae*. Selection and counterselection are the same as in panel C. Histones in this experiment were all expressed under the native *S*. *cerevisiae* histone promoters.

### *H. uvarum* H2A-H2B histone dimer is functionally divergent and incompatible with *S. cerevisiae*

Protein alignments showed that *Hanseniaspora* FEL histones diverged from the SEL and other yeasts, likely owing to the well-known rapid burst of evolution in the stem of the FEL (Figures S2&3; Steenwyk *et al*. 2019). To examine the functional significance of this divergence at the protein level, we used a histone replacement assay in *S. cerevisiae* (Haase *et al*. 2019). Using an *S*. *cerevisiae* strain in which the native histone clusters are deleted from their chromosomal loci and a single set of core histones genes is provided on a counter-selectable “Superloser” plasmid (Haase *et al*. 2019). Using subsequent plasmid shuffling, the native core histones genes may be readily swapped for an incoming set of heterologous histones genes (Figure 1B). As histones H2A and H3 were the most incompatible between yeast and human (Truong and Boeke 2017), we first exchanged individual *Hanseniaspora* species’ H2A and H3 histones and found that these two individual histones readily functioned in *S. cerevisiae*, whereas histones from FEL species generally performed worse than those from SEL species (Figure S4A–B). We next attempted to swap in all four *H. uvarum*’s histone genes under the control of *S. cerevisiae* promoters, observing that they did not complement in *S*. *cerevisiae* (Figure 1C, Figure S4C–D). To determine which of *H*. *uvarum*’s histones or pairs of histones are inviable in *S*. *cerevisiae*, we first individually replaced each of *H. uvarum*’s histones with the homolog from *S. cerevisiae*. We observed weak complementation when we replaced either *Huva*H2A or *Huva*H2B with *Sc*H2A or *Sc*H2B, respectively (Figure S4C–D), suggesting that the *Huva*H2A-H2B dimer is incompatible with *S. cerevisiae*. We confirmed this by replacing the *Huva*H2A-H2B dimer with the *Sc*H2A-H2B dimer (in the context of the *Huva*H3-H4 dimer), resulting in full complementation (Figure 1D). These complementation experiments neatly correlate with the divergence levels observed in the *Hanseniaspora* histones, where protein divergence increases from the lowest in H4 and H3 to the most divergent in H2A and H2B (Figure S2–4). Interestingly, this contrasts a previous report of human histone complementation in *S. cerevisiae*, where human H4 and H2B complemented very well, and human H3 and H2A did not complement (Truong and Boeke 2017). Moreover, the potent genetic suppressor of human histones in yeasts, *DAD1*^E50D^ (Haase *et al*. 2023a), did not rescue the inviability of *H*. *uvarum*’s H2A-H2B dimer (data not shown), suggesting that there may be species-specific incompatibilities at play.

### Core histone gene cis-regulatory innovation in *Hanseniaspora* FEL

A striking dominant negative effect was observed when the histone gene’s promoters of *H. uvarum* (*Huva*PRO) were used to express histones in *S*. *cerevisiae* (Figure 1C; column 4 or 6). Additionally, the *Huva*PRO was only viable when expressing *S. cerevisiae* histones, albeit with significantly reduced growth compared to native *S. cerevisiae* histone promoters (Figure 1C). We probed whether underlying sequence differences in these promoters decreased fitness. To this end, we examined the histone gene promoters (defined as the intervening sequence between the two divergently transcribed histone genes) from a set of *Hanseniaspora* and outgroup species and performed motif enrichments. The histone promoters varied in size, with FEL species and other Saccharomycotina yeasts having markedly shorter promoters than the SEL species, suggesting that histone promoter size might have increased prior to cis-regulatory divergence (Figure S5A). We identified a motif corresponding to the DNA binding sequence of the conserved core histone gene regulator Spt10 in most outgroup species, as expected, and throughout the SEL lineage (Figure 2A–B). The Spt10 motif was notably absent from species in the *Hanseniaspora* FEL lineage. From a discriminative *de novo* motif search, we identified a second motif corresponding to the DNA binding sequence of the transcription factor Mcm1 (Figure 2A–B). Mapping the presence of the Mcm1-like motif to the phylogeny showed that it was exclusive to histone promoters of the FEL lineage – suggesting that the FEL lineage underwent an ancestral histone gene regulatory rewiring event and that these sites may be responsible for *Huva*PRO reducing fitness in *S*. *cerevisiae*. We also identified a second motif that was enriched in the FEL species’ histone promoters corresponding to the Rap1 DNA binding site. However, this site was not found across all species in the FEL and was notably absent from the majority of the histone H3-H4 promoters; as such, we did not investigate this putative site further.

### Repeated innovations to core histone gene cis-regulatory mechanisms in yeasts

Additionally, we failed to identify the conserved Spt10 motif in histone promoters of the representative species from three basal genera: *Lipomyces*, *Tortispora*, and *Stramerella* (Figure 2A). Due to *Stramerella*’s phylogenetic placement, it is likely that the Spt10 motif was secondarily replaced; therefore, we focused on understanding cis-regulatory landscapes for the two outgroup lineages, *Lipomyces* and *Tortispora*/*Trigonopsis*. We searched for enrichment of histone promoter cis-regulatory motifs in additional species from these two lineages. In the most basal group, *Lipomyces*, we observed a complex cis-regulatory landscape for its core histone genes, marked by four significantly enriched GC-rich DNA motifs, for which we failed to find any significant similarity to known fungal DNA binding proteins (Figure S5C–D). In contrast, in the *Tortispora*/*Trigonopsis* lineage, we observed a strong enrichment for a cis-regulatory motif corresponding to the DNA binding motif of the Rfx1 transcription factor (Figure S5D). Taken together, these results suggest multiple rewiring events of histone cis-regulatory mechanisms early in the evolution of budding yeasts, though revealing the precise mechanisms used in these basal lineages will require further investigation.

**Figure 2.**
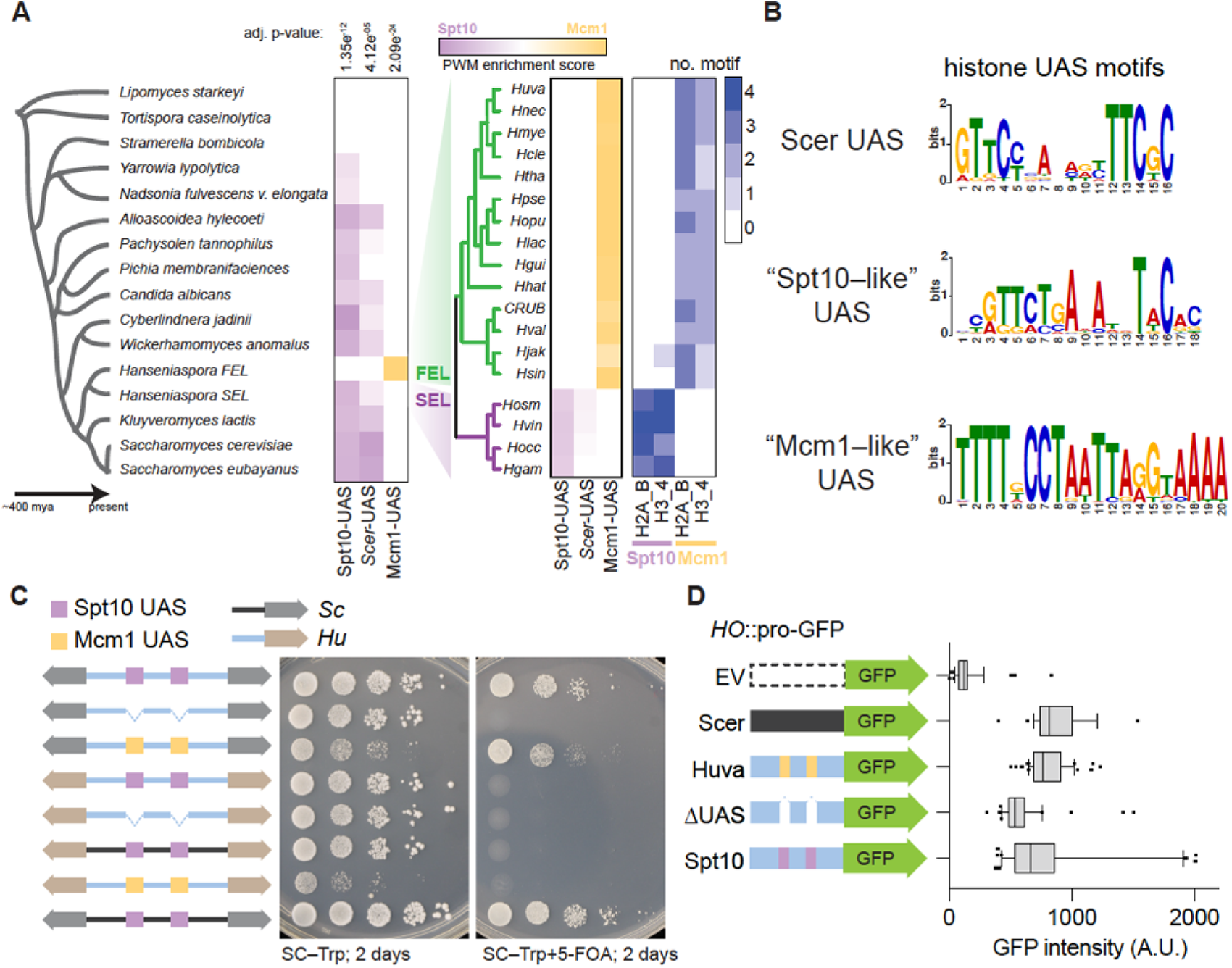
Core histone cis-regulatory rewiring in *Hanseniaspora* FEL. **A**. Histone promoter motif enrichment analysis of the Spt10 and Mcm1 DNA binding sites. Histone gene promoters were searched for enrichment of either *S*. *cerevisiae*’s Spt10 motif (Scer UAS), the conserved Spt10 motif found across taxa (“Spt10-like” UAS), and the Mcm1 motif found in *Hanseniapsora* FEL (“Mcm1-like” UAS) using the AME motif enrichment (MEME suite). The average position weight matrix (PWM) of all identified motifs from each species is shown, with a dual–color scheme showing enrichment for Spt10 or Mcm1 (purple to yellow). Lastly, the number of identified motifs is shown for the *Hanseniaspora* species. **B**. Motifs discovered (MEME) from a search of histone promoters from *Hanseniaspora* and four outgroup species (*S*. *cerevisiae*, *K*. *lactis*, *C. jadinii*, *W. anomalus*). The “Scer UAS” motif was constructed using known Spt10 DNA binding sites of the four core histones genes (Eriksson *et al*. 2012). **C**. Functional dissection of the Mcm1 binding sites in histone promoters of *H*. *uvarum*. Plasmid shuffle assay was carried out as in Figure 1B. Shown to the left of the growth assays are diagrammatic representations of the constructs tested. All four histones were swapped in all cases, but only one histone gene cluster is shown for simplicity (For H2A-H2B, three Mcm1 sites are present, and for H3-H4, two sites are present). Colored links between gene arrows represent the species of origin of the promoter used, whereas colored boxes represent the specific DNA binding element present at the UAS sites. Additionally, the species’ histones are color-coded, as shown. **D**. Expression analysis of a GFP. A GFP reporter targeting the *HO* locus with no upstream promoter or the indicated promoter (all HTA2 promoters) was transformed into BY4741. Total GFP fluorescence was measured from 10 images acquired using an EVOS M7000 imaging system.

### Mcm1 DNA binding sites underlie toxicity o*f H. uvarum* histone promoters in *S. cerevisiae*

Next, to determine whether the Mcm1 DNA binding sites were responsible for the growth defect, we generated versions *Huva*Pro with the Mcm1 sites removed (*Huva*Pro^ΔMcm1^). Deletion of Mcm1 sites resulted in amelioration of the toxic phenotype, and, as expected, the *Huva*Pro^ΔMcm1^ was no longer viable when expressing *S. cerevisiae* histones (Figure 2C). We restored viability to the *Huva*Pro^ΔMcm1^ construct by replacing the Mcm1 sites for the consensus Spt10 binding motif from the histone promoters of *S. cerevisiae (Huva*Pro^UAS-Spt10)^. The *Huva*Pro^UAS-Spt10^ was no longer toxic when driving the expression of the native *S. cerevisiae* histones (Figure 2C, SC–Trp), and, importantly, *Huva*Pro^UAS-Spt10^ was sufficient for viability when expressing the histones of *S. cerevisiae* (Figure 2B, SC–Trp+5-FOA). We conclude that the Mcm1 sites are the functional elements responsible for the toxicity in *S. cerevisiae*.

The toxicity of the histones could be due to overexpression, temporal misexpression, or both. We used a GFP reporter assay to investigate the level of activity from the promoters of H2A genes. We observed that the *Huva*Pro-H2A showed similar levels of GFP intensity to the native *Sc*Pro-H2A (Figure 2D; the *HTA1* promoter). Moreover, removing the Mcm1 UAS sites in *Huva*Pro-H2A reduced the expression of GFP, and the insertion of the Spt10 UAS restored expression to normal levels (Figure 2D). These data support the idea that the *H*. *uvarum* promoters do not lead to histone overexpression *per se* but perhaps temporal misexpression; although we did not formally test this, it potentially explains the striking dominant negative growth effect in *S*. *cerevisiae*, as expression of histones outside of S-phase has a well-known cytotoxic effect (Kurat *et al*. 2014). In conclusion, we show that the Mcm1-mode is incompatible in species with the ancestral Spt10-mode of histone gene regulation.

### *H. uvarum* Mcm1 functionally replaces *S. cerevisiae* Mcm1

The above data strongly indicate that Mcm1 regulates the core histones in *Hanseniaspora* FEL. To gain insight into whether the core histones shifted into a novel regulation paradigm or Mcm1 functionally diverged during the evolution of *Hanseniaspora* FEL, we developed two tests. First, we explored the evolution of cis-regulatory sites of Mcm1 target genes in *H. uvarum*, by performing a motif search of the Mcm1 DNA binding sequence in the promoters of orthologs of *Sc*Mcm1-regulated genes. We found that the large majority of orthologs of *Sc*Mcm1-regulated genes in *H. uvarum* also have Mcm1 binding sites in their putative promoter regions (Figure S6A), suggesting that *Huva*Mcm1 regulates the same set of target genes in *H. uvarum*.

**Figure 3.**
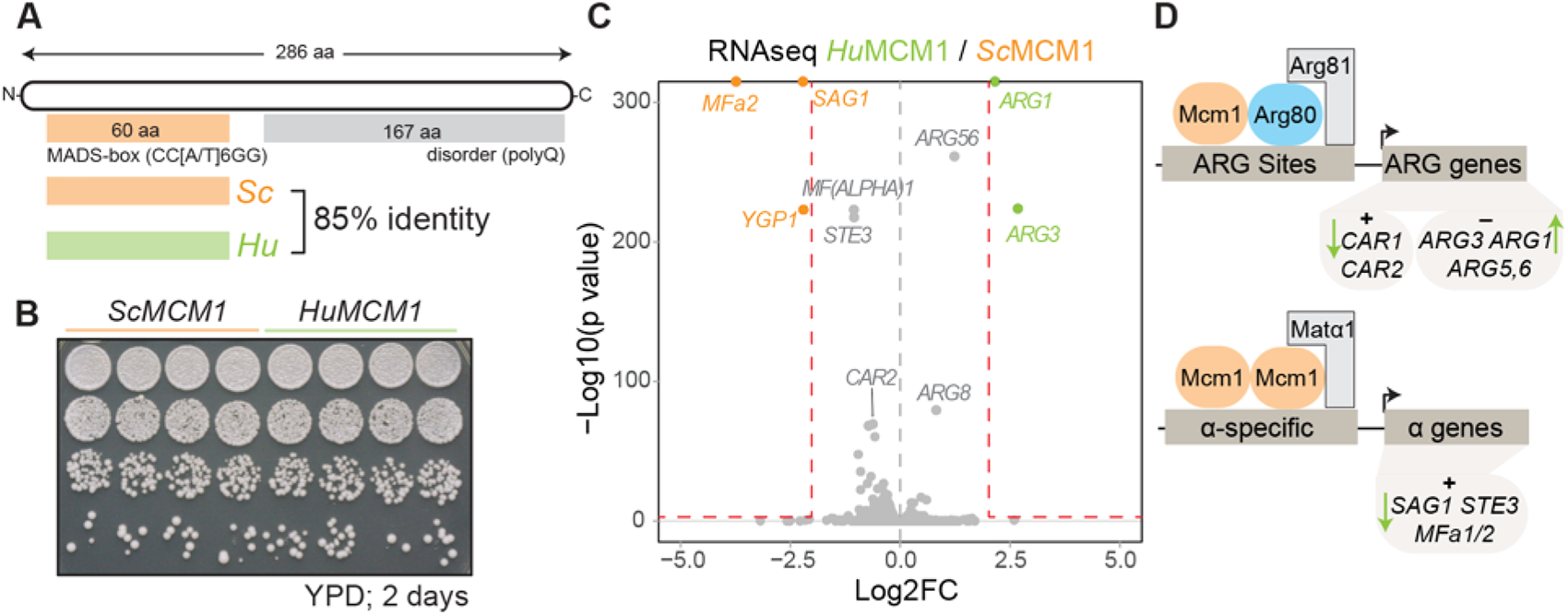
Mcm1 essential function and ancestral regulatory network are conserved. **A**. Diagram of the Mcm1 protein, highlighting the MADS-box (<u>M</u>CM1, <u>A</u>GAMOUS, <u>D</u>EFICIENS, <u>S</u>RF) domain, Mcm1’s essential DNA binding domain, and conservation between *S*. *cerevisiae* (*Sc*Mcm1; orange) and *H*. *uvarum* (*HuvaMcm1* green). The MADS-box domain of Mcm1 from *H*. *uvarum* directly replaced *S*. *cerevisiae*’s Mcm1 MADS-box domain. **B–C**. *HuvaMcm1* has little effect on the *S*. *cerevisiae* transcriptome. (**B**) Fitness assay of WT and the HuvaMcm1 *S*. *cerevisiae* strains. Strains carrying *Sc*Mcm1 or *HuvaMcm1* were grown overnight in YPD medium; the following day, cultures were normalized (A_600_ ≈ 1.0), and cells were spotted on agar plates and grown for two days at 30°C. (**C**) Volcano plot comparing log2 fold change in gene expression between yeast with native *Sc*Mcm1 and strains with *HuvaMcm1* . Significantl dysregulated genes (log2FC>2 or <-2 and p-value < 0.01) are colored depending on the direction of change; upregulated in *HuvaMcm1*, green; and downregulated in *HuvaMcm1*, orange. Genes regulated by Arg80/Mcm1 are highlighted. Related to supplemental figures 8 and 9. **D**. *HuvaMcm1* behaves like the pre-duplication *Anc*MADS (computationally inferred ancestor of Mcm1 and Arg80 (Baker *et al*. 2013)), as Mcm1 activated ARG genes (*CAR1*, *CAR2*) have lower expression, Mcm1 repressed genes (*ARG3*, *ARG1*, *ARG5,6*) are upregulated, and α–specific genes (Mcm1-activated; *SAG1*, *STE3*, *MFa1/2*) show reduced expression in the strain with *HuvaMcm1* . Green arrows show the direction of expression change in the *HuvaMcm1* strains.

Next, we genetically tested *Huva*Mcm1 function in *S*. *cerevisiae* by examining whether *HuvaMcm1* complements Mcm1 in *S*. *cerevisiae* (Figure 3A). Specifically, we tested for the essential function of Mcm1, which is conferred by the DNA-binding MADS-box domain (Figure 3A, Figure S7A). We were able to generate Mcm1::*HuvaMcm1* MADS-box domain replacements via CRISPR-Cas9 genome editing at the native locus, demonstrating that *HuvaMcm1* retains the essential functions of *Sc*Mcm1 (Figure S7A–C). Additionally, these strains showed no phenotypic difference (Figure 3B). Assessment of the transcriptomic effects of *HuvaMcm1* replacement by RNA sequencing showed that *HuvaMcm1* had little effect on *S. cerevisiae*’s transcriptome, with only five genes being significantly dysregulated (Figure 3C; Figure S7D–E). *Huva*Mcm1 represents a living ancestor of the *MCM1* gene prior to the segmental duplication that gave rise to the paralogous gene *ARG80* (Messenguy and Dubois 1993). Prior work on a computationally inferred Mcm1/Arg80 ancestral gene (*Anc*MADS; note the previously reconstructed ancestral Mcm1/ARG80 was positioned at a divergence time of ∼92 MYA; in comparison, *HuvaMcm1* represents a divergence time of ∼138 MYA, suggesting much deeper conservation in function than previously indicated; divergence times taken from Shen *et al*. 2018) showed that *Anc*MADS led to dysregulation of MADS-Box activated ARG (arginine) genes, MADS-Box repressed ARG genes, MADS-Box activated mating genes, and MADS-Box repressed mating genes (Baker *et al*. 2013). In agreement, we observed dysregulation of this exact same set of genes, confirming that *HuvaMcm1* functions like the computationally determined *Anc*MADS (Figure 3C). Interestingly, deletion of the Mcm1 paralog, Arg80, sharpened toxicity of *Huva*Pro (Figure S6B), consistent with the idea that removal of competitive binding by Arg80 potentially increases binding of Mcm1 to *Huva*Pro and thus potentiates toxic effects of histone misexpression caused by *Huva*Pro. These data support the conclusion that *HuvaMcm1* is conserved in function and is consistent with a model in which the core histones were rewired into a preexisting regulatory network rather than *HuvaMcm1* diverging in function in *Hanseniaspora* FEL.

### Rapid cell division and decoupled core histone and DNA synthesis *in H. uvarum*

Given the cis-regulatory divergence of core histone genes, we were curious whether the dynamics of core histone expression were altered in *H*. *uvarum*. Using a recently described system for the genetic manipulation of *H. uvarum,* we inserted an H2A–mNeonGreen (a LanYFP-derived fluorophore; Shaner *et al*. 2013) fusion construct at the native *HTA1* locus (HTA-mNG; Figure 4A–B). Similar systems have been used to monitor cell cycle dynamics and histone protein synthesis in *S*. *cerevisiae* (Garmendia-Torres *et al*. 2018). We tracked the nuclear intensity of H2A over the course of a few cell cycles (Figure 4C–F), measuring an average cell cycle length of ∼60 minutes for cells grown in SC without exposure to the excitation laser (Figure S8A; Video S1). In contrast, when cells were grown with exposure every 5 minutes, we observed they had a slightly increased cell cycle length of ∼80 minutes, which we observed in either WT or fused H2A-mNG cells, suggesting a slight phototoxic effect (Figure S8A; Video S2). The cell cycle dynamics in *H. uvarum* are markedly different than in *S. cerevisiae* in several ways (for comparative data from *S. cerevisiae*, we refer the reader to Garmendia-Torres *et al*. 2018). First, both daughter and mother cells synchronously bud following cytokinesis (Figure 4C; Video S1), whereas in *S*. *cerevisiae,* daughter cells display prolonged G1 and S phases. Second, histone synthesis begins well after bud emergence (∼30 minutes; Figure 4E–F), in contrast to *S*. *cerevisiae,* where histone synthesis begins just prior to bud emergence. Third, the nascent bud reaches near–full mature cell size just prior to mother-daughter cytokinesis (Figure 4D; Video S2); however, in *S*. *cerevisiae,* the nascent bud does not reach the size of the mother until later cell cycles. Taken together, the non-canonical features of the *H. uvarum* cell cycle may promote rapid cell division.

**Figure 4.**
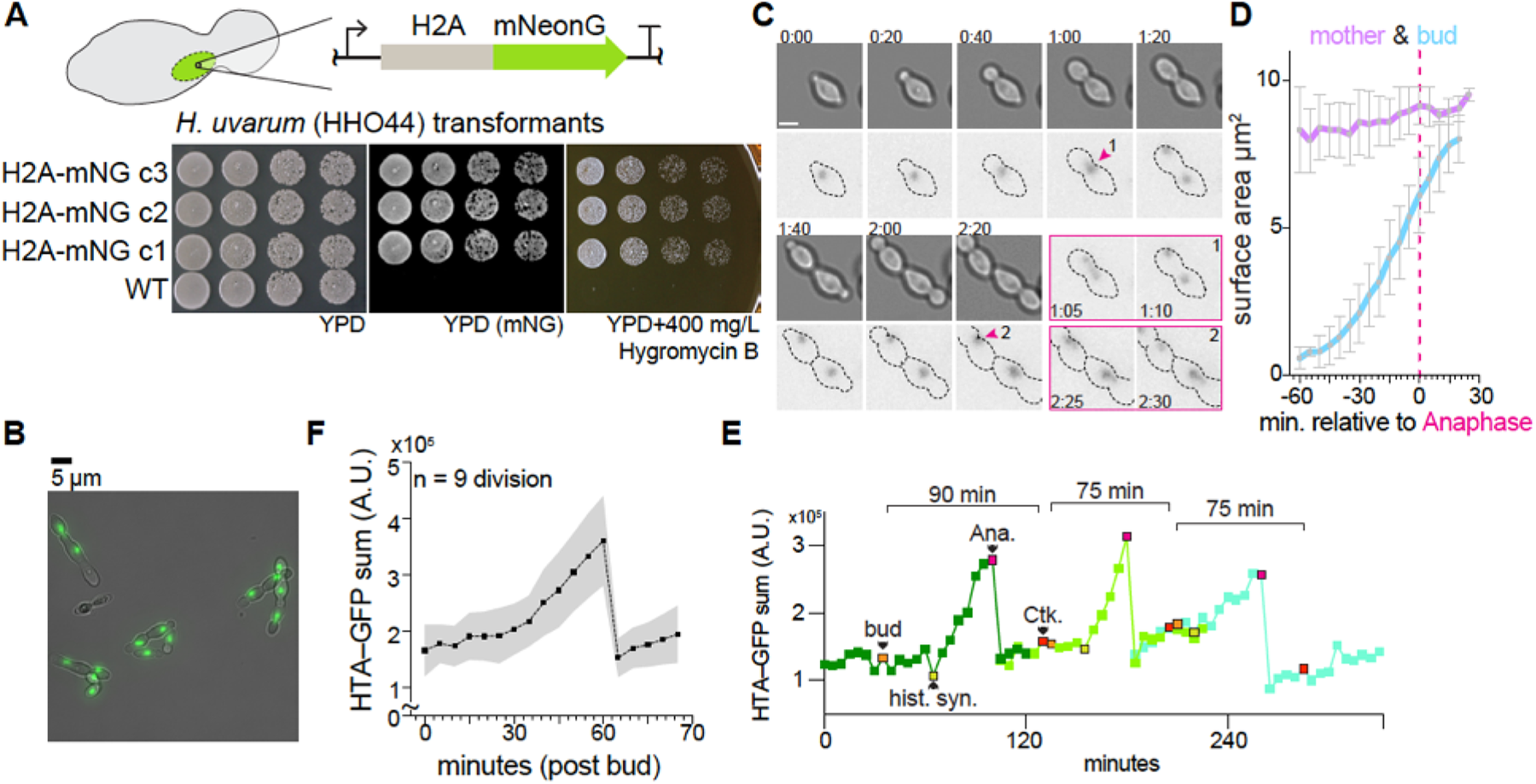
Cell cycle dynamics and histone synthesis in *H*. *uvarum*. **A**. Schematic of H2A-mNeonGreen-hphMX cassette used to target the endogenous H2A gene. Note that the hphMX cassette used for hygromycin B resistance is not depicted but is immediately downstream of the terminator of the H2A-mNG fusion. Below is a spot assay of three transformed clones grown on YPD (imaged on a Bio-Rad ChemiDoc MP imaging system with blue epi-illumination) and YPD with 400 mg/L Hygromycin B. **B**. Example image of *H*. *uvarum* cells with H2A-mNG tag. Scale bar = 5 µm. **C**. Time-lapse growth of *H*. *uvarum* with H2A-mNG tag. A series of images are shown from Video S2 at intervals of twenty minutes. Two divisions are separately shown in the images outlined in magenta. Scale bar = 2.5 µm; time H:MM. **D**. Mother and daughter (bud) cell surface area analysis. The surface area was determined by manually outlining mother and daughter pairs (n = 5). Mean areas with ±SD are shown. The bud grew ∼8-fold faster than the mother cell; mother = 0.84 ±0.24 µm^2^ h^-1^; bud = 6.6 ±0.3 µm^2^ h^-1^. **E**. Example track of H2A-mNG levels from time-lapse growth of *H*. *uvarum* cells. Three divisions are followed, an orange-colored point indicates bud emergence (bud); a yellow point indicates the start of histone synthesis (hist. syn.); a magenta point indicates the start of mitosis/anaphase (Ana.); a red point indicates the completion of division (cytokinesis; Ck.). Time-lapse images were acquired every 5 minutes at 30°C in SC medium. **F**. Average H2A-mNG levels during the cell cycle of *H*. *uvarum*. Movies from nine cells were all set relative to bud emergence (t =0), and H2A levels were tracked and quantified until cytokinesis.

In most eukaryotes, histone protein synthesis is tightly coupled to the status of DNA replication (Eriksson *et al*. 2012; Rattray and Müller 2012; Kurat *et al*. 2014). In *S*. *cerevisiae,* histone synthesis is inhibited following a DNA replication block with the treatment of hydroxyurea (HU) via a specific regulatory coupling (Bhagwat *et al*. 2021), which is likely mediated by the HIR complex (<u>hi</u>stone regulatory genes (*HIR1*, *HIR2*, *HIR3*) and <u>h</u>istone <u>p</u>eriodic <u>c</u>ontrol gene (*HPC2*)). Given the altered cell cycle dynamics, we were curious whether histone synthesis was dependent on DNA synthesis in *H. uvarum*. We arrested cells in early S-phase using HU and followed cell division after release (Video S3). We observed that after 60 minutes of HU arrest, histone levels were near their expected mitotic levels (Figure 5A–E), whereas the DNA remained unreplicated, as confirmed by DNA content analysis by flow cytometry (Figure 5B). Following HU release, the cells restarted and completed DNA synthesis within ∼60 minutes (Figure 5B); however, during this same period, histone levels remained constant (Figure 5C). We next tracked H2A-mNG levels in single cells following HU release. We confirmed that histone levels do not increase in individual cells during the first cell division after HU release (Figure 5D–G), with most cells completing division within ∼60–120 minutes after HU release (Figure S8B). Moreover, we observed that in the subsequent division, histones were normally synthesized, and the total histone levels reached a significantly higher maximal level than during the first division following HU arrest (Figure 5D–G). Intriguingly, the mother cells divided significantly slower in this second division than their corresponding daughter cell (Figure S8C). In sum, histone synthesis in *H. uvarum* is independent of status of DNA synthesis, as histone levels were at mitotic levels prior to completion of DNA synthesis and did not increase during DNA replication.

Intriguingly, the regulatory decoupling of histone and DNA synthesis in *H*. *uvarum* is similar to the behavior of *hir* mutants in *S*. *cerevisiae,* in which hydroxyurea-mediated repression of histone gene expression likely depends on direct recruitment of the HIR complex to the histone promoter via the N-terminal region of Hpc2. Remarkably, *HPC2* was lost ancestrally in the *Hanseniaspora* FEL (Steenwyk *et al*. 2019), suggesting that the loss of this negative regulation modality, perhaps alongside the shift to Mcm1 regulation, enabled loss of direct coupling between histone and DNA synthesis. The significance of the “Mcm1-mode” remains to be clarified *in vivo* for *H*. *uvarum*, especially in the context of gene loss of key cell cycle regulators (Figure 5F). In conclusion, monitoring the dynamics of histone levels in live cells across the cell cycle has revealed that histone synthesis in *H*. *uvarum* is decoupled from DNA synthesis, and we speculate that the regulatory rewiring of its histone genes into the Mcm1– mode, together with the loss of Hpc2 and critical cell cycle regulators, were potentially adaptive for the emergence of its rapid cell cycle progression.

**Figure 5.**
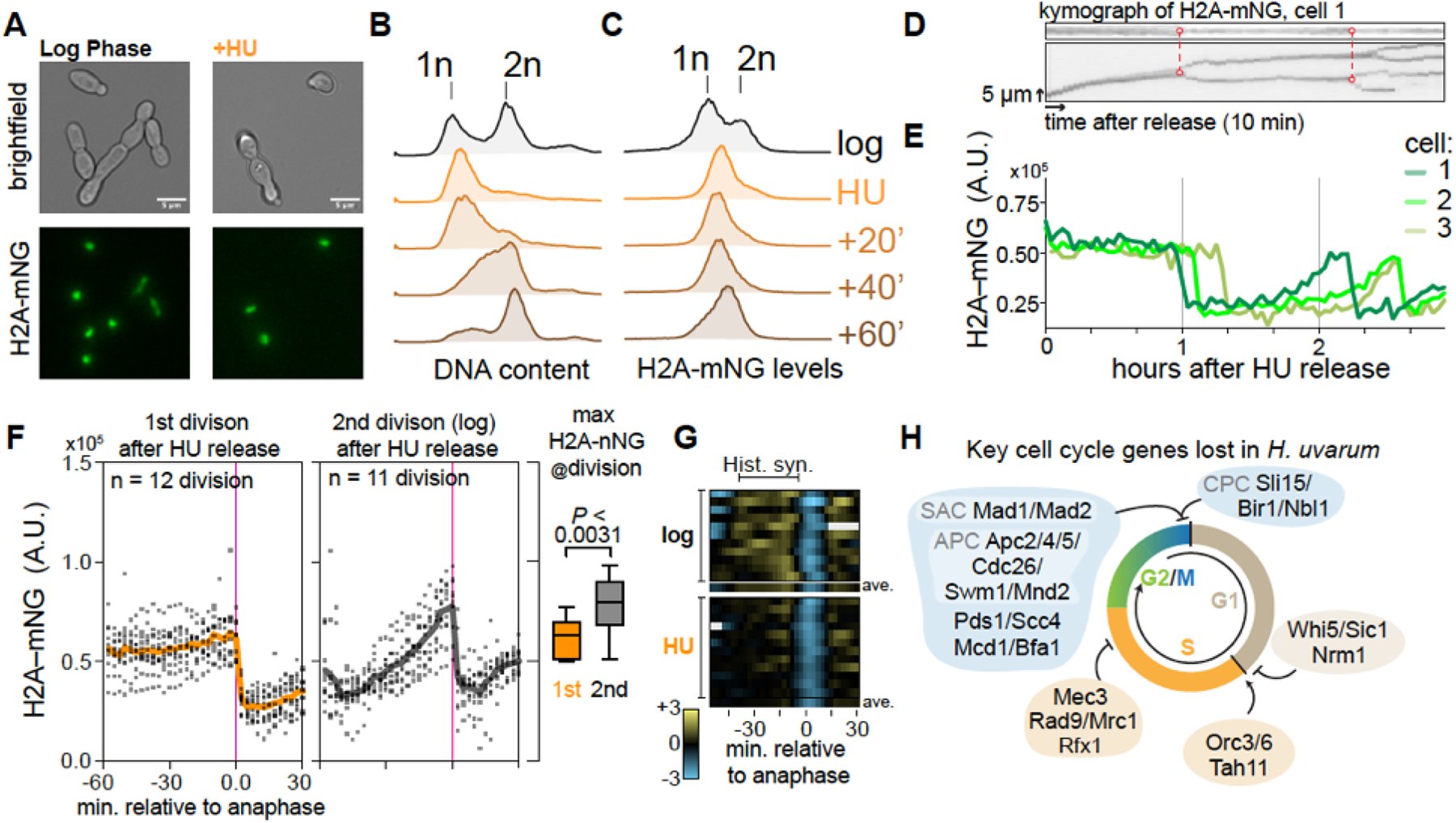
Core histone synthesis is decoupled from DNA synthesis in *H*. *uvarum*. **A**. Example images of cells from mid-log phase growth or after 60 minutes of arrest in 300 mM HU. **B–C**. DNA content analysis and histone abundance (H2A-mNG level) by flow cytometry of cells from mid-log phase growth or after, and following release, from 60 minutes of arrest in HU. **D–E**. Histone levels do not appreciably increase in single cells following release from HU. (**D**) Kymographs of H2A-mNG levels from cell 1 (the top kymograph is focused on the nucleus of the mother cell, and the bottom is a kymograph focused on a region containing the nuclei of mother and daughter cells), connected red dots indicating the time of division, and dashed lines denote the two divisions tracked below. The arrow representing time corresponds to 10 minutes, and the arrow representing space corresponds to 5 µm. (**E**) Example tracks of the sum of H2A-mNG levels (arbitrary units; A.U.) in three mother cells monitored for up to ∼3 hours post-release from HU. **F**. Histone level dynamics in the first and second cell division after HU arrest. H2A-mNG levels were quantified relative to anaphase (magenta line, timepoint = 0), at which time histone levels remained constant prior to the first division after release from HU. In contrast, histone levels showed a characteristic linear increase during the second division after HU release. Average profiles are shown as solid lines. To the right is the maximum H2A-mNG signal just prior to anaphase, showing that histone levels were higher during the second cell division after HU release. **G**. Rate of histone synthesis (H2A-mNG) in cells after HU arrest (n = 12) or during the second round of division (log; n = 11). The first derivative of H2A-mNG levels is plotted for each replicate cell. Positive values (yellow) correspond to instantaneous increases in histone levels (synthesis), no change corresponds to zero (black), and negative values (blue) correspond to instantaneous decreases in histone levels (anaphase). Missing values are filled in as gray rectangles. Each rectangle represents a 5-minute interval. Below each condition, the average profile of that condition is plotted. **H**. Overview of key cell cycle regulators lost in *H*. *uvarum*. Data from Steenwyk *et al*. 2019. We hypothesize that a common signal initiates DNA replication and histone synthesis in *H*. *uvarum*; however, due to the loss of negative regulation (loss of *HPC2*) and the shift to the Mcm1-mode of positive regulation, core histone synthesis is no longer coupled to the status of DNA replication. As various checkpoints during the cell cycle have been lost, it may be advantageous to fully commit to histone production even if DNA replication is perturbed. Thus, while gene loss and cis-regulatory rewiring potentially led to the loss of the DNA synthesis-dependent regulation of histone synthesis, *H*. *uvarum* ensures histones are produced in a timely manner consistent with its rapid progression through the cell cycle.

## Conclusions

We have detailed the evolution of core histone gene clusters across the budding yeast phylogeny (Figure 6). We can infer that the last budding yeast common ancestor (BYCA) had a single copy of each core histone gene cluster, where each histone gene was interspaced with at least two introns, and ancestral transcriptional regulation was likely carried out by yet-to-be-identified trans–regulators which bind GC-rich motifs present in these BYCA promoter sequences. The emergence and fixation of the Spt10 regulatory mode is likely ancient (∼320-380 MYA) and occurred after the divergence of the Trigonopsidaceae, but before the divergence of the Dipodascaceae/Trichomonascaceae, as the extant species in *Tortispora* and *Trigonopsis* regulate their core histones with a cis–motif predicted to bind to the Rfx1 transcription factor, suggesting that in the earliest lineages, there was strong diversifying selection on the mode of core histone gene regulation from the ancestral mode seen in *Lipomyces*. Based on the presence and number of histone gene clusters, we predict that the histone gene clusters duplicated twice independently after the emergence of the Spt10 regulatory paradigm (once in the ancestor of *Nadsonia* and once in the ancestor of the species *Alloascoidea hylecoeti* and *S. cerevisiae*). Thus, the contemporary set of core histone gene clusters and regulatory mode (Spt10) emerged after the divergence of the Alloascoideaceae (∼250 MYA).

Intriguingly, one set of core histone genes was lost alongside a cis-regulatory rewiring event we identified in *Hanseniaspora*. As the events occurred in the stem of the FEL we cannot determine the precise timing of these events, although gene loss occurring prior to rewiring would allow for fewer needed cis-regulatory changes to fix the Mcm1-mode. Additionally, the SEL species *H*. *gamundiae* may represent a case of convergent evolution at a more intermediate state, where partial histone paralogous gene loss occurred without cis-regulatory changes. In addition to the association of gene loss and cis-regulatory rewiring, we also observed that histone gene duplications occurred alongside predicted cis-regulatory changes in more basal lineages. Together, these observations suggest that, in general, copy number changes are associated with cis-regulatory changes of the core histones, although the broader significance of such an association remains to be investigated in other lineages.

**Figure 6.**
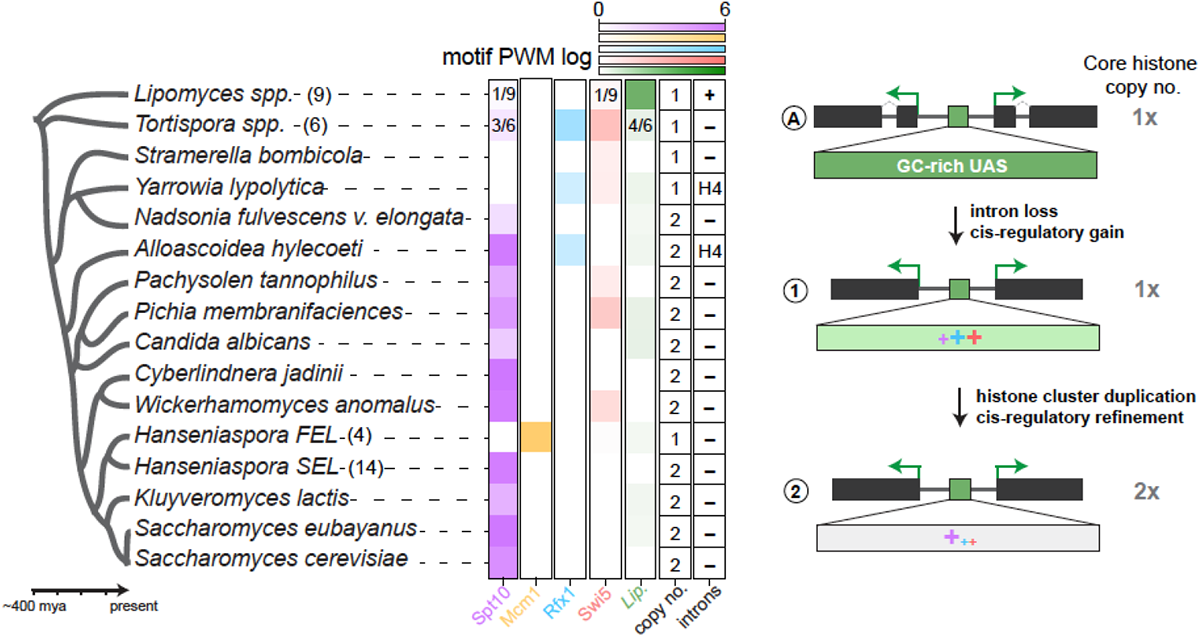
A model for evolution of core histone genes in Saccharomycotina. Overview of motif enrichment analysis of core histone promoters and the broad strokes of core histone evolution. For lineages for which we examined multiple species, the number of species is given in parenthesis; moreover, if only some species showed enrichment, we show the fraction of species with each motif in their histone promoters. We outline three major transitions that likely gave rise to the current state of core histones in yeasts. First, introns were lost from the ancestral histone genes, and various modes of cis-regulation emerged from a GC-rich UAS ancestor. Second, the histone clusters duplicated and various lineages refined their core histone cis-regulation (the majority fixed the Spt10 mode, whereas some lineages, such as *Tortispora*, fixed the Rfx1/Swi5 mode). Cladogram is based on the phylogeny from Shen *et al*. 2018.

Lastly, we uncovered an innovation to core histone gene regulation in the *Hanseniaspora* FEL, where an Mcm1-mode replaced the ancestral Spt10-mode. We speculate that this led to the altered cell-cycle regulation of its core histone genes, potentially alongside the loss of *HPC2,* decoupling histone synthesis from the status of DNA synthesis. Further work will be needed to uncover the temporal dynamics of DNA replication and other key cell cycle events at the single-cell level in *Hanseniaspora*. Given the uniqueness of genes absent from the *Hanseniaspora* FEL, we believe it can be a future model for cell biology for understanding how typically essential processes function in the absence of key cell cycle regulators. Moreover, framing future studies into an evolutionary cell biology framework (Helsen *et al*. 2023) will enhance future studies into the biochemical and molecular details of core histone gene regulation in *Hanseniaspora* and budding yeast more broadly.

## Contributions

M.A.B.H. conceptualized the project, performed the formal investigation, wrote the manuscript, and prepared figures. J.L.S. provided comments and suggestions on the manuscript. J.D.B. supervised the research and provided funding. All authors edited the manuscript.

## Acknowledgments

We thank Luciana Lazar-Stefanita for her helpful comments and support; Jürgen Heinisch for kindly providing strains, plasmids, and the transformation protocol for *Hanseniapsora uvarum*. We thank Gregory Goldberg for helpful comments on early drafts. We are grateful to Chris Hittinger and Antonis Rokas and the members of their laboratories for fruitful discussions of this work. An NSF Rules of Life grant supported this work: Epigenetics 2 (award number: MCB1921641) to J.D.B. J.L.S. is a Howard Hughes Medical Institute Awardee of the Life Sciences Research Foundation.

## Data availability

All yeast strains and plasmids are available upon request to Jef D. Boeke. Raw RNA sequencing data related to Figure 3 were deposited to Sequence Read Archive (SRA) and are available under the Bioproject ID PRJNA987614.

## Methods

### Yeast strains and plasmids

All strains and plasmids are listed in Supplemental Tables 1 and 2, respectively. Both *S*. *cerevisiae* and *H*. *uvarum* strains were grown in standard yeast media (YPD or SC) at 30°C. *S*. *cerevisiae* strains were transformed using standard lithium acetate procedures (Gietz and Schiestl 2007). We transformed *H*. *uvarum* by electroporation following a previously published method with modifications (Heinisch *et al*. 2023). Briefly, a 50 mL overnight culture (A_600_ ≈ 0.1) was grown at 30°C for 16 hours at 210 rpm. The next day the density of the culture was checked; if A_600_ ≈ 3.5–4.0, we immediately proceeded to the following steps; however, if A_600_ >4.0, we diluted the culture and let it grow for an appropriate amount of time. Cells were then collected, washed 1x in sterile water, resuspended in Lithium acetate buffer (10 mM Tris, pH 8.0, 1 mM EDTA, 100 mM Lithium acetate, 10 mM DTT), and incubated for 1 hour at room temperature with agitation. Next cells were collected, washed in ice-cold sterile water, and then washed 2x in ice-cold 1M sorbitol. Finally, cells were collected and resuspended in 500 µL of ice-cold 1M sorbitol (this mixture can be frozen at –80°C for later transformation). For one transformation, we used an aliquot of 100 µL of the cell suspension. A maximum of 10 µL of DNA (∼1 µg) was placed into a 2 mm cuvette, and cells were added onto and mixed by lightly tapping the cuvette; each cuvette was then incubated at room temperature for 10 minutes. Next, samples were electroporated using a Bio-Rad MicroPulser with the “SC2” default settings. After electroporation, the cell suspension was diluted with 1 mL fresh YPD medium, transferred to a 1.5 mL centrifuge tube, and incubated with rotation for 3–4 hours. Lastly, cells were collected, and the transformation was plated to YPD + 400 µg of hygromycin B.

### Histone gene presence and absence in Hanseniaspora

We initially searched a set of published *Hanseniaspora* genomes and four outgroup species for histone genes with BLASTP, using the protein sequences of *S*. *cerevisiae*’s histones as the query. Each hit was manually inspected and verified to remove erroneous hits such as histone variants H2A.Z or CenH3 (Cse4). For species of the FEL, we determined which of the histone clusters were lost through synteny analysis, comparing *Hanseniaspora* genomes to *S*. *cerevisiae* and the inferred pre-WGD ancestor using the Yeast Gene Order Browser.

### Dual plasmid histone shuffling and plasmid cloning

We modified our plasmid shuffling tool set for histone humanization in order to shuffle in the *Hanseniaspora* histone genes and promoters. Briefly, a histone shuffle strain (yMAH302), which has all eight chromosomally encoded core histone genes deleted and a single copy of each core histone encoded on a counter-selectable plasmid, is transformed with an incoming set of histone genes. By counter-selection of the *URA3* marker with 5–FOA, the cells are forced to use the incoming set of histone genes (*TRP1* marker), and the assay readout is cell growth. We used the low-background ‘Superloser’ shuffle plasmid to reduce the frequency of spontaneous *ura3* mutants that lead to erroneous 5-FOA^R^ colonies (Haase *et al*. 2019). For our histone shuffling assays, we used the native histone DNA sequences from *Hanseniaspora* species, as we observed that codon-optimized versions complemented worse (data not shown). We followed a general workflow for plasmid shuffle assay as previously published (Haase *et al*. 2019, 2023a).

### Core histone gene promoter analysis

Core histone promoters were defined as the intervening sequence between the divergently transcribed histone genes. Regulatory motif enrichment analysis was done as previously described with the following variations (Haase *et al*. 2021). Using AME (Analysis of Motif Enrichment, default options), we searched each histone promoter for the canonical Spt10 UAS from *S*. *cerevisiae*. Next, using MEME (maximum width = 24, site distribution = anr), we identified the two motifs, ‘Spt10-like’ and ‘Mcm1-like’, from a search of all histone promoter sequences. We then again used the AME search (now using either the ‘Spt10-like’ and ‘Mcm1-like’ motif) to determine which species each motif was associated with, observing the strong enrichment of the ‘Mcm1-like’ motif in the *Hanseniaspora* FEL. We performed similar motif enrichment analysis for targeted searches of the outgroup lineages *Lipomyces* and *Tortispora*/*Trigonopsis*.

### Histone promoter UAS replacement assays

We precisely deleted, or replaced with a consensus Spt10 UAS, the putative Mcm1 binding sites from the *Huva*PRO (both H2A-H2B and H3-H4). These constructs were then used for plasmid shuffle assays to assess their function. For measurements of GFP expression, we used a ubiquitin-N-degron GFP reporter (Houser *et al*. 2012) integrated at the HO locus with either no upstream promoter or the indicated promoter, as shown in Figure 2D. Total cellular fluorescence was determined from single cells from a series of images acquired on an EVOS M7000 imaging system.

### Mcm1 motif discovery of Mcm1-regulated orthologs in *H. uvarum*

We identified a list of Mcm1-regulated genes in *S. cerevisiae* and extracted the protein sequences of these genes. We then used BLASTP to search the genome of *H. uvarum* for orthologs of these Mcm1-regulated genes. Finally, we extracted 500 bp upstream from the start codon of each ortholog and searched for the Mcm1 DNA binding motif in each upstream region using the AME function of the MEME suite.

### Mcm1 replacement assay

We used CRISPR-Cas9 editing to directly replace the MADS-box domain of Mcm1 in *S*. *cerevisiae* (strain yMAH302). We edited Mcm1 MADS-box with a sgRNA (5’– AACGACTAGCAACAGGACCT–3’) targeting a Pam site overlapping codons 56 and 57, and the repair was directed using a dsDNA donor encoding the MADS-box domain of *Huva*Mcm1. Edited colonies were screened by diagnostic PCR/digestions, where the PCR-amplified Mcm1 MADS-box fragment from successfully edited clones was only positively digested with KpnI. Successfully edited clones were Sanger sequenced to confirm the edit. In addition, we isolated WT-edited clones, which only carried the sgRNA abolishing synonymous mutations, and used these as our control strains in the RNAseq experiment, ensuring any bias introduced via CRISPR-Cas9 cloning was correctly controlled.

We then grew strains to mid-log phase (A600 ∼0.6–0.8) and extracted RNA as previously reported (Haase *et al*. 2023a). We prepared total RNA with rRNA-depletion sequencing libraries using the QIAseq Stranded Total RNA kit (Qiagen Cat. 180745) and the QIAseq FastSelect-rRNA Yeast Kit (Qiagen Cat. 334217). Lastly, libraries were sequenced with an Illumina NextSeq 500 with paired-end 2 x 150 bp read chemistry.

### Flow cytometry and HU arrests

Cells were grown overnight in YPD at 30°C, and the following morning, saturated cultures were diluted to A_600_ ≈ 0.2 and grown until mid–log phase, A_600_ ≈ 0.6. Cells were then washed in PBS, resuspended in YPD + 300 mM hydroxyurea, and placed at room temperature for 60 minutes with agitation. Arrested cells were then washed 2x in fresh YPD and then resuspended in YPD, and placed at 30°C for outgrowth after HU arrests. We took aliquots of the cell suspension at various time points for analysis. For DNA content analysis, cells were first crosslinked with 0.5% paraformaldehyde for 15 minutes at 4°C. After 2x washes with PBS, crosslinked cells were then resuspended in ice-cold (-20°C) 70% methanol and incubated at 4°C for one hour. Cells were then washed with 2x PBS, resuspended in PBS + 2.5 µM SytoxGreen, and incubated at 30°C for 30 minutes. For analysis of HTA-mNeonGreen, aliquots of cells were taken at the appropriate time points, washed 2x in PBS, and placed on ice until analyzed for flow cytometry. Cells were then analyzed using a spectral cell analyzer (Sony SA3800), and data from approximately ∼30,000 events were analyzed in the FlowJo software.

### Time-lapse imaging of *H*. *uvarum*

Prior to imaging cells were grown to mid-log phase (A_600_ ≈ 0.6–0.8) in YPD. Cells were then collected, resuspended in SC medium, and placed at room temperature for one hour. Meanwhile, we prepared a 15 µ-slide VI (ibidi Cat. XXX) for imaging by coating the surface with Concanavalin A from *Canavalia ensiformis* (10 mg/mL in water). Cells were then loaded onto the slide and incubated for 10 minutes prior to two washing steps with SC media. Finally, cells were placed into a temperature-controlled EVOS M7000 imaging system, and time-lapses were collected at 30°C with images taken at either 2.5- or 5-minute intervals. For time-lapses after HU arrests, cells were arrested with HU as above. After HU arrest, cells were quickly washed in PBS and resuspended in SC medium, and immediately placed into the imaging chamber. Time lapses were then acquired the same as above. Movies were then analyzed in Fiji using the TrackMate plugin.

## Supplementary Figure Legends

**Supplementary Figure 1.** Paralogous gene loss of core histones in *Hanseniaspora*

**A.** Phylogeny of the *Hanseniaspora* and four outgroup species from Steenwyk *et al*. 2019, with the presence and absence of the core histones genes. Purple, the slow-evolving lineage; Green, the fast-evolving lineage.

**B–C**. Synteny analysis of the *HTA2B2* and *HTA1B1* gene clusters. Data for the pre-WGD ancestor and S. cerevisiae were taken from the yeast gene order browser (citation), and gene order was manually inferred for *H*. *vineae* (SEL) and *H*. *uvarum* (FEL). Similar results were obtained for the *HHT1F1* and *HHT2F2* gene clusters, in which *HHT1F1* was lost in the FEL (data not shown).

**Supplementary Figure 2.** Histone H3 and H4 protein alignments

Histone protein sequences were aligned using Mafft (v7) using “Auto” setting. Variants are color-coded as follows; red changes from *S. cerevisiae* histone sequence, blue residues altered in *S. cerevisiae* histone sequences.

Supplementary Figure 3. Histone H2A and H2B protein alignments

Histone protein sequences were aligned using Mafft (v7) using “Auto” setting. Variants are color-coded as follows; red changes from *S. cerevisiae* histone sequence, blue residues altered in *S. cerevisiae* histone sequences.

**Supplementary Figure 4.** H2A and H2B swaps

**A.** Histone swaps of H2A orthologs from SEL (*H*. *gamundiae*, *H*. *vineae*) and FEL (*H*. sp.CRUB/*valbyensis*, *H*. *hatyaiensis*, *H. uvarum*) species. Quantifications of 5FOA^R^ frequency are taken from three biological replicate swap experiments, with the mean and SD shown.

**B.** Histone swaps of H3 orthologs from SEL (*H*. *gamundiae*, *H*. *vineae*) and FEL (*H*. sp.CRUB/*valbyensis*, *H*. *hatyaiensis*, *H. uvarum*) species. Quantifications of 5FOA^R^ frequency are taken from three biological replicate swap experiments, with the mean and SD shown.

**C–D**. Histone swaps of *H*. *uvarum* core histone genes and with individual replacements with *S*. *cerevisiae* histones. (**C**) To identify which of the four histones from *H. uvarum* were inviable in *S*. *cerevisiae*, we replaced each single *H*. *uvarum* histone with its ortholog from *S*. *cerevisiae*. Only when we replaced either HuvaH2A or HuvaH2B did we observe weak complementation, indicating the H2A–H2B dimer combination is not viable. (**D**) The number of cells for each complementation was scaled up (plated ∼10^8^ cells) and plated onto an entire 10 cm petri dish.

**Supplementary Figure 5.** Diversity of core histone cis-regulatory mechanisms in basal lineages

**A.** Core histone promoter sizes in outgroup species, SEL, and FEL species.

**B–C**. Overview of histone gene structure and cis-regulation in the *Lipomyces*. Histone genes are at a single copy (except for H4, which has two copies), and each histone gene contains introns. Enrichment for DNA motifs from the core histone promoter revealed four significant motifs that are not present in other yeast species.

**D–E**. Overview of histone gene structure and cis-regulation in *Tortispora*/*Trigonopsis*. Histone genes are at a single copy and histone genes lack introns. Enrichment for DNA motifs from the core histone promoters revealed two significant motifs that are not present in other yeast species.

**Supplementar**y Figure 6. Mcm1 targets genes are conserved in *H*. *uvarum*

**A.** Forward search for Mcm1 binding sites in orthologs to *S*. *cerevisiae* Mcm1 regulated genes (M-to-G1) in two *Hanseniaspora* species.

**B.** Dot assays of *H*. *uvarum* histones and histone promoters in wildtype and Δ*arg80* strains.

**Supplementary Figure 7.** Editing the Mcm1 MADS-box domain

**A.** Primary sequence and secondary Mcm1 DNA binding domain structure from *H*. *uvarum* and *S*. *cerevisiae*. Modified from Messenguy and Dubois (2003).

**B.** Example transformation from the CRISPR-Cas9 editing. Left, the small guide RNA was transformed without a repair template; Right, sgRNA was transformed with a repair template, resulting in colony growth (sgRNA-resistant clones).

**C.** Genotyping of clones by PCR/digestions. PCR amplicons from the edited clones were digested with diagnostic enzymes EcoNI and KpnI. Successfully edited clones digest with only KpnI, whereas recombinants digest by both, and wildtype digests with EcoNI. Clones were confirmed by Sanger sequencing.

**D.** RNA extractions from wildtype (ScerMcm1) and mutant (HuvaMcm1) strains.

**E.** Transcriptional changes to genes involved in arginine metabolism due to loss of Mcm1 repression (*ARG1*, *ARG3*, *ARG5,6*, and *ARG8*). Log2FC is shown as a green-colored box for each gene, with intensity increasing with upregulation. Additionally, the z-score expression value of each gene is given for the two conditions; left, ScerMcm1; right, HuvaMcm1.

**Supplementary Figure 8.** Cell cycle length in *H*. *uvarum*

**A.** Doubling time inferred from time-lapse movies of *H. uvarum* grown with and without (red x) exposure to the excitation laser.

**B.** Time to cell cycle completion after release from HU arrest, tracking the mother cell and nascent daughter cell. A single mother cell is shown in the timestamped image sequence to the left, and the daughter cell completes its cell cycle ∼20 minutes prior to its mother. Right is data from 14 mother cells and the subsequent divisions.

**Video S1.** *H*. *uvarum* H2A-mNG tagged strain growth.

Cells from a mid-log phase culture (A_600_ ≈ 0.6) were secured to the chamber surface and imaged every 2.5 minutes at 30°C. Only phase contrast images were taken. Scale bar 5 µM; time HH:MM:SS.

**Video S2.** *H*. *uvarum* H2A-mNG tagged strain growth with GFP excitation.

Cells from a mid-log phase culture (A_600_ ≈ 0.6) were secured to the chamber surface and imaged every 5 minutes at 30°C. Images were acquired from both the RFP channel (mNeonGreen is a LanYFP-derived fluorophore, as such, it is excited by the RFP laser (531 nm), which we found to be less phototoxic than the GFP laser (470 nm)) and phase contrast. Scale bar 10 µM; time HH:MM:SS.

**Video S3.** *H*. *uvarum* H2A-mNG tagged strain growth with GFP excitation after HU release. Cells were first arrested in HU for 45 minutes and washed 2x in fresh SC medium prior to being secured to the chamber surface. We mixed an equal proportion of untagged and H2A-mNG-tagged cells prior to imaging. Time-lapse images were acquired every 5 minutes at 30°C. Scale bar 25 µM; time HH:MM:SS.

## References

Badura J., N. Van Wyk, S. Brezina, I. S. Pretorius, D. Rauhut, et al., 2021 Development of Genetic Modification Tools for Hanseniasporauvarum. IJMS 22: 1943. 10.3390/ijms22041943

Badura J., N. van Wyk, K. Zimmer, I. S. Pretorius, C. von Wallbrunn, et al., 2023 PCR-based gene targeting in Hanseniaspora uvarum. FEMS Yeast Research 23: foad034. 10.1093/femsyr/foad034

Baker C. R., V. Hanson-Smith, and A. D. Johnson, 2013 Following Gene Duplication, Paralog Interference Constrains Transcriptional Circuit Evolution. Science 342: 104–108. 10.1126/science.1240810

Becher P. G., A. Hagman, V. Verschut, A. Chakraborty, E. Rozpędowska, et al., 2018 Chemical signaling and insect attraction is a conserved trait in yeasts. Ecol Evol 8: 2962–2974. 10.1002/ece3.3905

Bhagwat M., S. Nagar, P. Kaur, R. Mehta, I. Vancurova, et al., 2021 Replication stress inhibits synthesis of histone mRNAs in yeast by removing Spt10p and Spt21p from the histone promoters. Journal of Biological Chemistry 297: 101246. 10.1016/j.jbc.2021.101246

Dollard C., S. L. Ricupero-Hovasse, G. Natsoulis, J. D. Boeke, and F. Winston, 1994 SPT10 and SPT21 are required for transcription of particular histone genes in Saccharomyces cerevisiae. Mol Cell Biol 14: 5223–5228. 10.1128/mcb.14.8.5223-5228.1994

Eriksson P. R., G. Mendiratta, N. B. McLaughlin, T. G. Wolfsberg, L. Mariño-Ramírez, et al., 2005 Global regulation by the yeast Spt10 protein is mediated through chromatin structure and the histone upstream activating sequence elements. Mol Cell Biol 25: 9127–9137. 10.1128/MCB.25.20.9127-9137.2005

Eriksson P. R., D. Ganguli, V. Nagarajavel, and D. J. Clark, 2012 Regulation of Histone Gene Expression in Budding Yeast. Genetics 191: 7–20. 10.1534/genetics.112.140145

Garmendia-Torres C., O. Tassy, A. Matifas, N. Molina, and G. Charvin, 2018 Multiple inputs ensure yeast cell size homeostasis during cell cycle progression. eLife 7: e34025. 10.7554/eLife.34025

Gietz R. D., and R. H. Schiestl, 2007 High-efficiency yeast transformation using the LiAc/SS carrier DNA/PEG method. Nat Protoc 2: 31–34. 10.1038/nprot.2007.13

Goffeau A., B. G. Barrell, H. Bussey, R. W. Davis, B. Dujon, et al., 1996 Life with 6000 Genes. Science 274: 546–567. 10.1126/science.274.5287.546

Groenewald M., C. T. Hittinger, K. Bensch, D. A. Opulente, X.-X. Shen, et al., 2023 A genome-informed higher rank classification of the biotechnologically important fungal subphylum Saccharomycotina. Studies in Mycology 105: 1–22. 10.3114/sim.2023.105.01

Haase M. A. B., D. M. Truong, and J. D. Boeke, 2019 Superloser: A Plasmid Shuffling Vector for Saccharomyces cerevisiae with Exceedingly Low Background. G3: Genes, Genomes, Genetics 9: 2699–2707. 10.1534/g3.119.400325

Haase M. A. B., J. Kominek, D. A. Opulente, X.-X. Shen, A. L. LaBella, et al., 2021 Repeated horizontal gene transfer of GAL actose metabolism genes violates Dollo’s law of irreversible loss. Genetics 217: iyaa012. 10.1093/genetics/iyaa012

Haase M. A. B., G. Ólafsson, R. L. Flores, E. Boakye-Ansah, A. Zelter, et al., 2023a DASH /Dam1 complex mutants stabilize ploidy in histone-humanized yeast by weakening kinetochore-microtubule attachments. The EMBO Journal. 10.15252/embj.2022112600

Haase M. A. B., L. Lazar-Stefanita, G. Ólafsson, A. Wudzinska, M. J. Shen, et al., 2023b Human macroH2A1 drives nucleosome dephasing and genome instability in histone-humanized yeast. bioRxiv. 10.1101/2023.05.06.538725

Hamby K. A., A. Hernández, K. Boundy-Mills, and F. G. Zalom, 2012 Associations of Yeasts with Spotted-Wing Drosophila (Drosophila suzukii; Diptera: Drosophilidae) in Cherries and Raspberries. Appl Environ Microbiol 78: 4869–4873. 10.1128/AEM.00841-12

Hauer M. H., A. Seeber, V. Singh, R. Thierry, R. Sack, et al., 2017 Histone degradation in response to DNA damage enhances chromatin dynamics and recombination rates. Nat Struct Mol Biol 24: 99–107. 10.1038/nsmb.3347

Heinisch J. J., A. Murra, K. Jürgens, and H.-P. Schmitz, 2023 A Versatile Toolset for Genetic Manipulation of the Wine Yeast Hanseniaspora uvarum. IJMS 24: 1859. 10.3390/ijms24031859

Helsen J., G. Sherlock, and G. Dey, 2023 Experimental evolution for cell biology. Trends in Cell Biology S0962892423000806. 10.1016/j.tcb.2023.04.006

Houser J. R., E. Ford, S. M. Chatterjea, S. Maleri, T. C. Elston, et al., 2012 An improved short-lived fluorescent protein transcriptional reporter for Saccharomyces cerevisiae. Yeast 29: 519–530. 10.1002/yea.2932

Jensen L. J., T. S. Jensen, U. de Lichtenberg, S. Brunak, and P. Bork, 2006 Co-evolution of transcriptional and post-translational cell-cycle regulation. Nature 443: 594–597. 10.1038/nature05186

Kurat C. F., J. Recht, E. Radovani, T. Durbic, B. Andrews, et al., 2014 Regulation of histone gene transcription in yeast. Cell. Mol. Life Sci. 71: 599–613. 10.1007/s00018-013-1443-9

Lazar-Stefanita L., M. A. B. Haase, and J. D. Boeke, 2023 Humanized nucleosomes reshape replication initiation and rDNA/nucleolar integrity in yeast. bioRxiv. 10.1101/2023.05.06.539710

Luger K., A. W. Mäder, R. K. Richmond, D. F. Sargent, and T. J. Richmond, 1997 Crystal structure of the nucleosome core particle at 2.8 Å resolution. Nature 389: 251–260. 10.1038/38444

MacAlpine D. M., and G. Almouzni, 2013 Chromatin and DNA replication. Cold Spring Harb Perspect Biol 5: a010207. 10.1101/cshperspect.a010207

Malik H. S., and S. Henikoff, 2003 Phylogenomics of the nucleosome. Nat Struct Mol Biol 10: 882–891. 10.1038/nsb996

Mariño-Ramírez L., I. K. Jordan, and D. Landsman, 2006 Multiple independent evolutionary solutions to core histone gene regulation. Genome Biol 7: R122. 10.1186/gb-2006-7-12-r122

Messenguy F., and E. Dubois, 1993 Genetic Evidence for a Role for MCM1 in the Regulation of Arginine Metabolism in Saccharomyces cerevisiae. Molecular and Cellular Biology 13: 2586–2592. 10.1128/mcb.13.4.2586-2592.1993

Messenguy F., and E. Dubois, 2003 Role of MADS box proteins and their cofactors in combinatorial control of gene expression and cell development. Gene 316: 1–21. 10.1016/S0378-1119(03)00747-9

Osley M. A., 1991 THE REGULATION OF HISTONE SYNTHESIS IN THE CELL CYCLE. Annu. Rev. Biochem. 60: 827–861. 10.1146/annurev.bi.60.070191.004143

Rattray A. M. J., and B. Müller, 2012 The control of histone gene expression. Biochemical Society Transactions 40: 880–885. 10.1042/BST20120065

Robbins E., and T. W. Borun, 1967 THE CYTOPLASMIC SYNTHESIS OF HISTONES IN HELA CELLS AND ITS TEMPORAL RELATIONSHIP TO DNA REPLICATION. Proc. Natl. Acad. Sci. U.S.A. 57: 409–416. 10.1073/pnas.57.2.409

Rueda-Mejia M. P., R. A. Ortiz-Merino, S. Lutz, C. H. Ahrens, M. Künzler, et al., 2022 Pantothenate auxotrophy in a naturally occurring biocontrol yeast. bioRxiv. 10.1101/2022.12.14.519733

Saubin M., H. Devillers, L. Proust, C. Brier, C. Grondin, et al., 2020 Investigation of Genetic Relationships Between Hanseniaspora Species Found in Grape Musts Revealed Interspecific Hybrids With Dynamic Genome Structures. Front. Microbiol. 10: 2960. 10.3389/fmicb.2019.02960

Schwarz L. V., M. J. Valera, A. P. L. Delamare, F. Carrau, and S. Echeverrigaray, 2022 A peculiar cell cycle arrest at g2/m stage during the stationary phase of growth in the wine yeast Hanseniaspora vineae. Current Research in Microbial Sciences 3: 100129. 10.1016/j.crmicr.2022.100129

Shaner N. C., G. G. Lambert, A. Chammas, Y. Ni, P. J. Cranfill, et al., 2013 A bright monomeric green fluorescent protein derived from Branchiostoma lanceolatum. Nat Methods 10: 407–409. 10.1038/nmeth.2413

Shen X.-X., D. A. Opulente, J. Kominek, X. Zhou, J. L. Steenwyk, et al., 2018 Tempo and Mode of Genome Evolution in the Budding Yeast Subphylum. Cell 175: 1533–1545.e20. 10.1016/j.cell.2018.10.023

Steensels J., and K. J. Verstrepen, 2014 Taming Wild Yeast: Potential of Conventional and Nonconventional Yeasts in Industrial Fermentations. Annu. Rev. Microbiol. 68: 61–80. 10.1146/annurev-micro-091213-113025

Steenwyk J. L., D. A. Opulente, J. Kominek, X.-X. Shen, X. Zhou, et al., 2019 Extensive loss of cell-cycle and DNA repair genes in an ancient lineage of bipolar budding yeasts. PLOS Biology 17: e3000255. 10.1371/journal.pbio.3000255

Truong D. M., and J. D. Boeke, 2017 Resetting the Yeast Epigenome with Human Nucleosomes. Cell 171: 1508–1519.e13. 10.1016/j.cell.2017.10.043

Van Wyk N., J. Badura, C. Von Wallbrunn, and I. S. Pretorius, 2023 Exploring future applications of the apiculate yeast Hanseniaspora. Critical Reviews in Biotechnology 1–20. 10.1080/07388551.2022.2136565

Venkatesh S., and J. L. Workman, 2015 Histone exchange, chromatin structure and the regulation of transcription. Nat Rev Mol Cell Biol 16: 178–189. 10.1038/nrm3941

Wei Y., L. Yu, J. Bowen, M. A. Gorovsky, and C. D. Allis, 1999 Phosphorylation of histone H3 is required for proper chromosome condensation and segregation. Cell 97: 99–109. 10.1016/s0092-8674(00)80718-7

Yun C.-S., and H. Nishida, 2011 Distribution of Introns in Fungal Histone Genes, (J. Stajich, Ed.). PLoS ONE 6: e16548. 10.1371/journal.pone.0016548

