## Supplemental Figures for "Gene loss and cis-regulatory novelty shaped core histone gene evolution in the apiculate yeast *Hanseniaspora uvarum*"

Supplementary Figure 1. Paralogous gene loss of core histones in *Hanseniaspora*

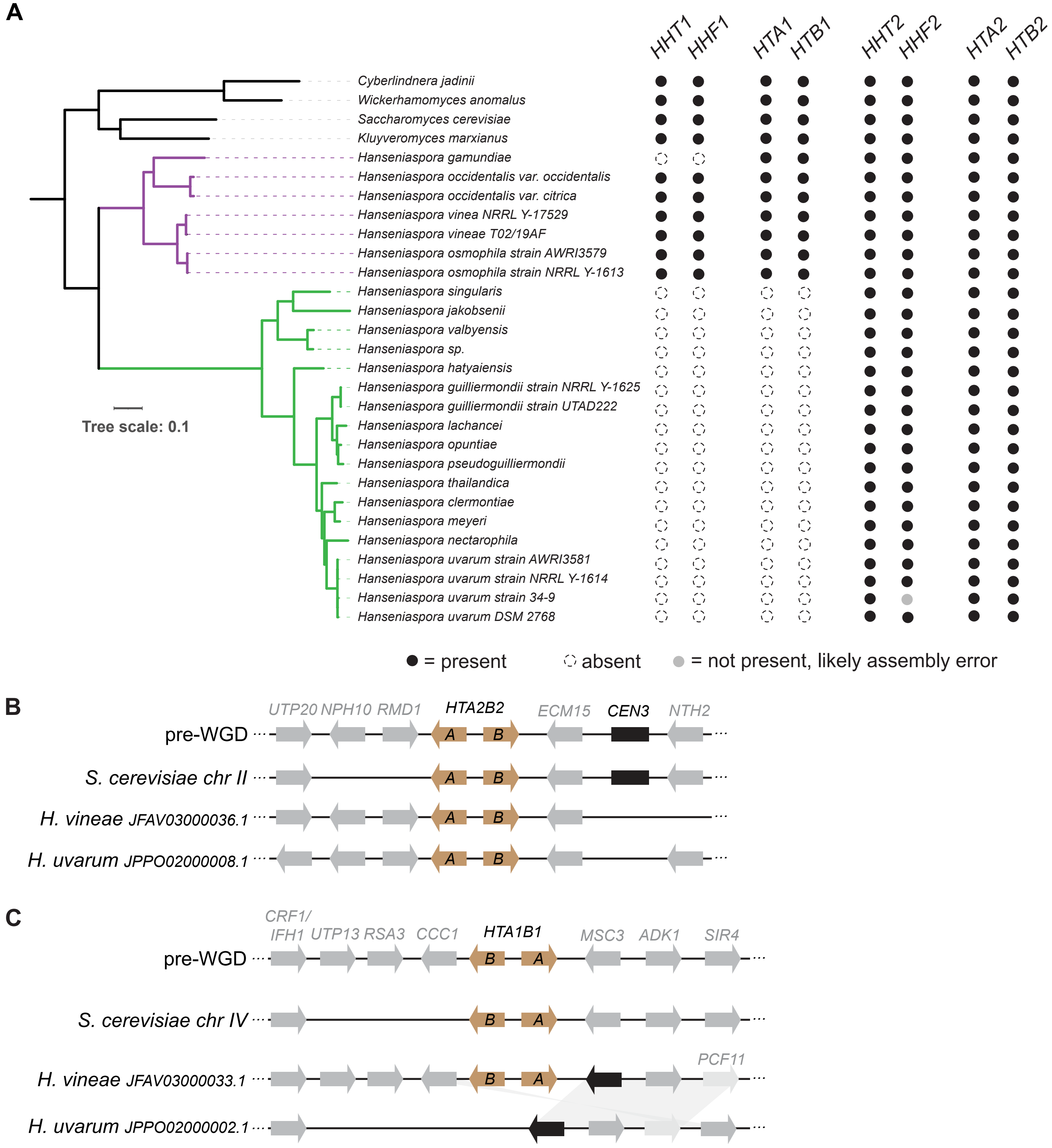

### Supplementary Figure 2. Histone H3 and H4 protein alignments

#### Histone H4

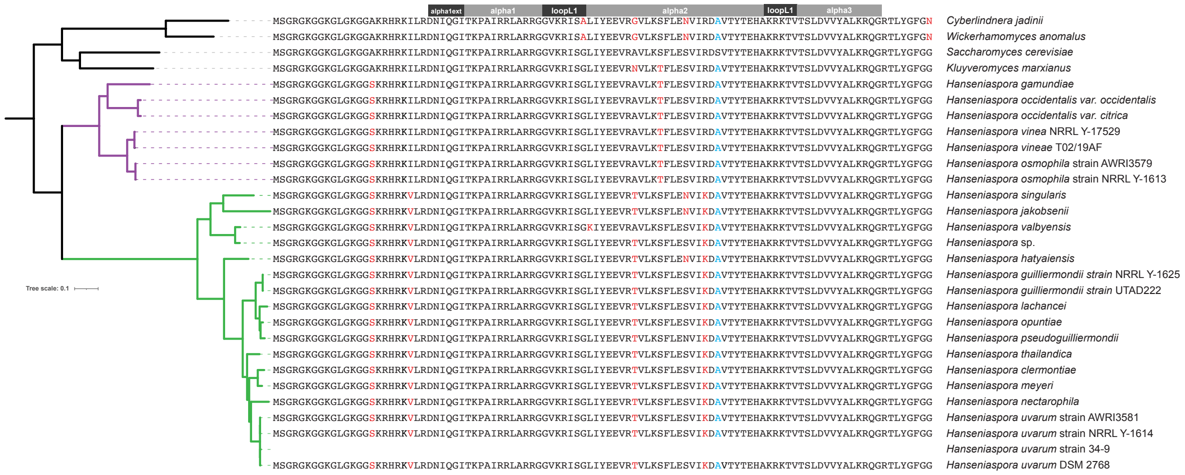


#### Histone H3

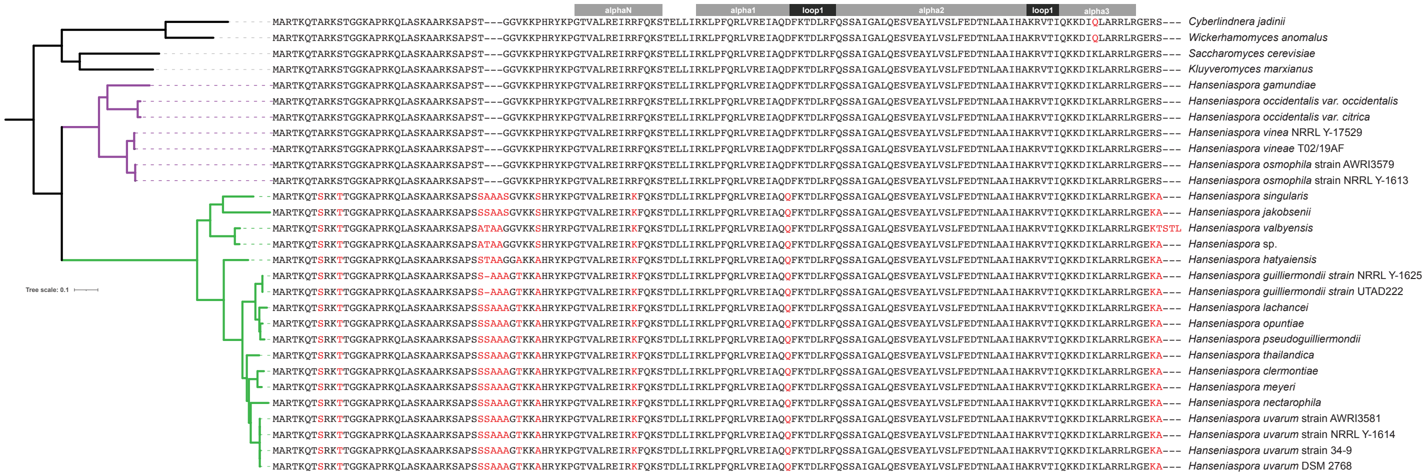

### Supplementary Figure 3. Histone H2A and H2B protein alignments

#### Histone H2A

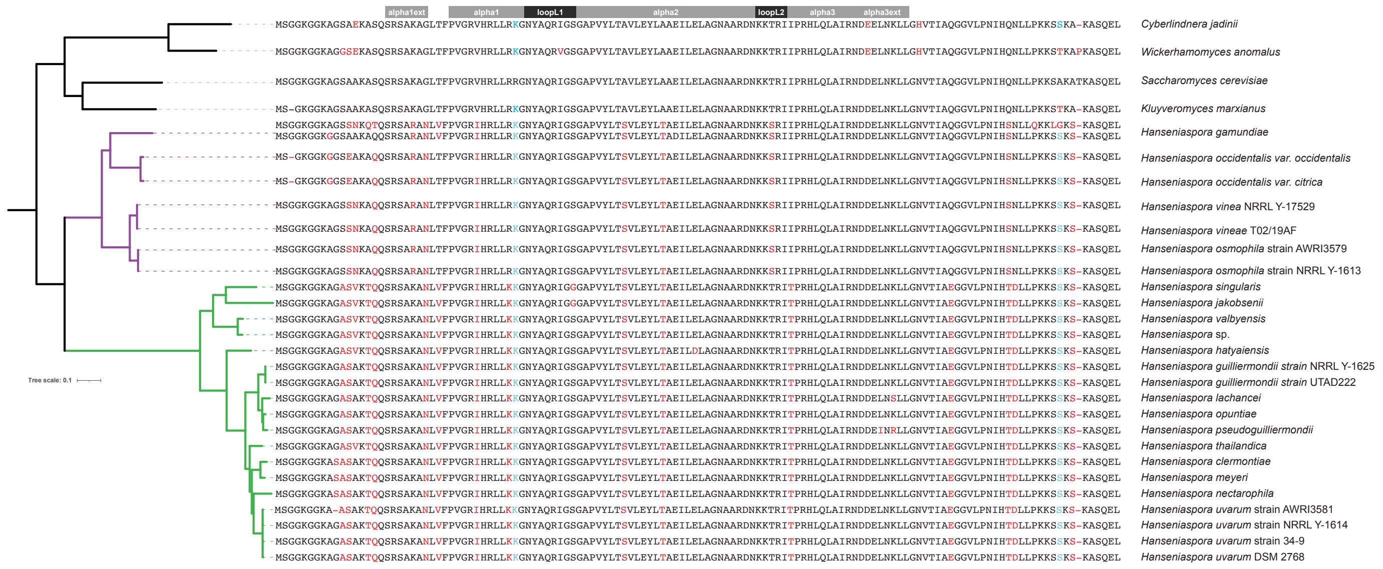


#### Histone H2B

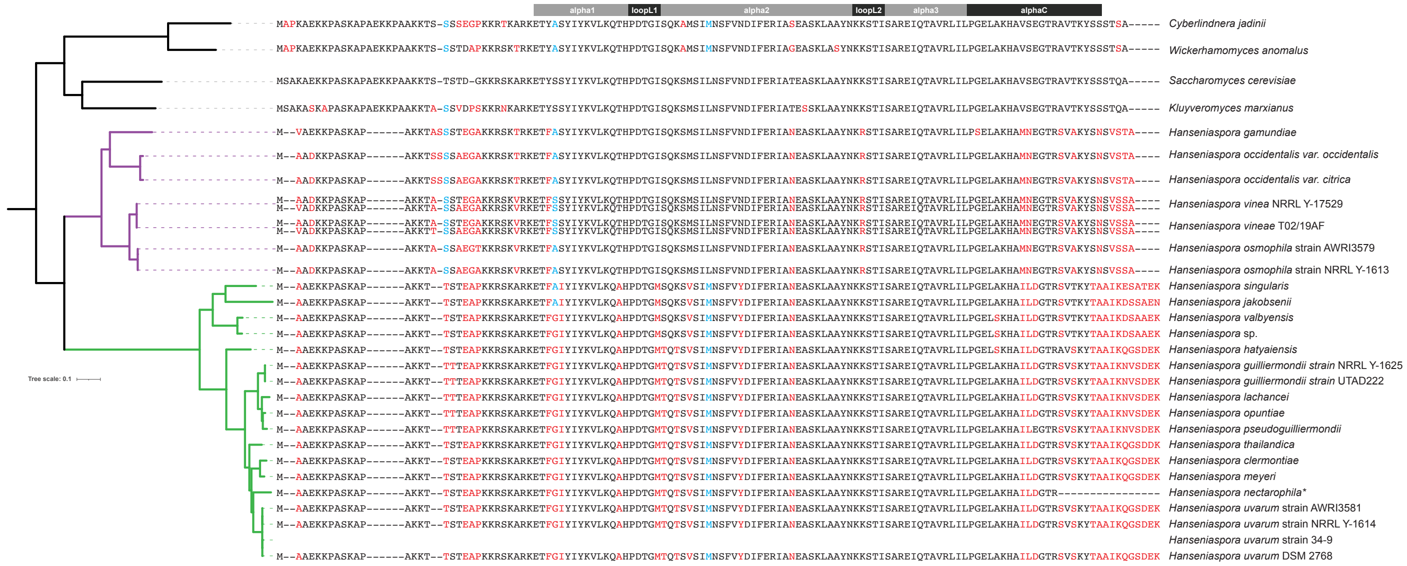

Supplementary Figure 4. H2A and H2B swaps

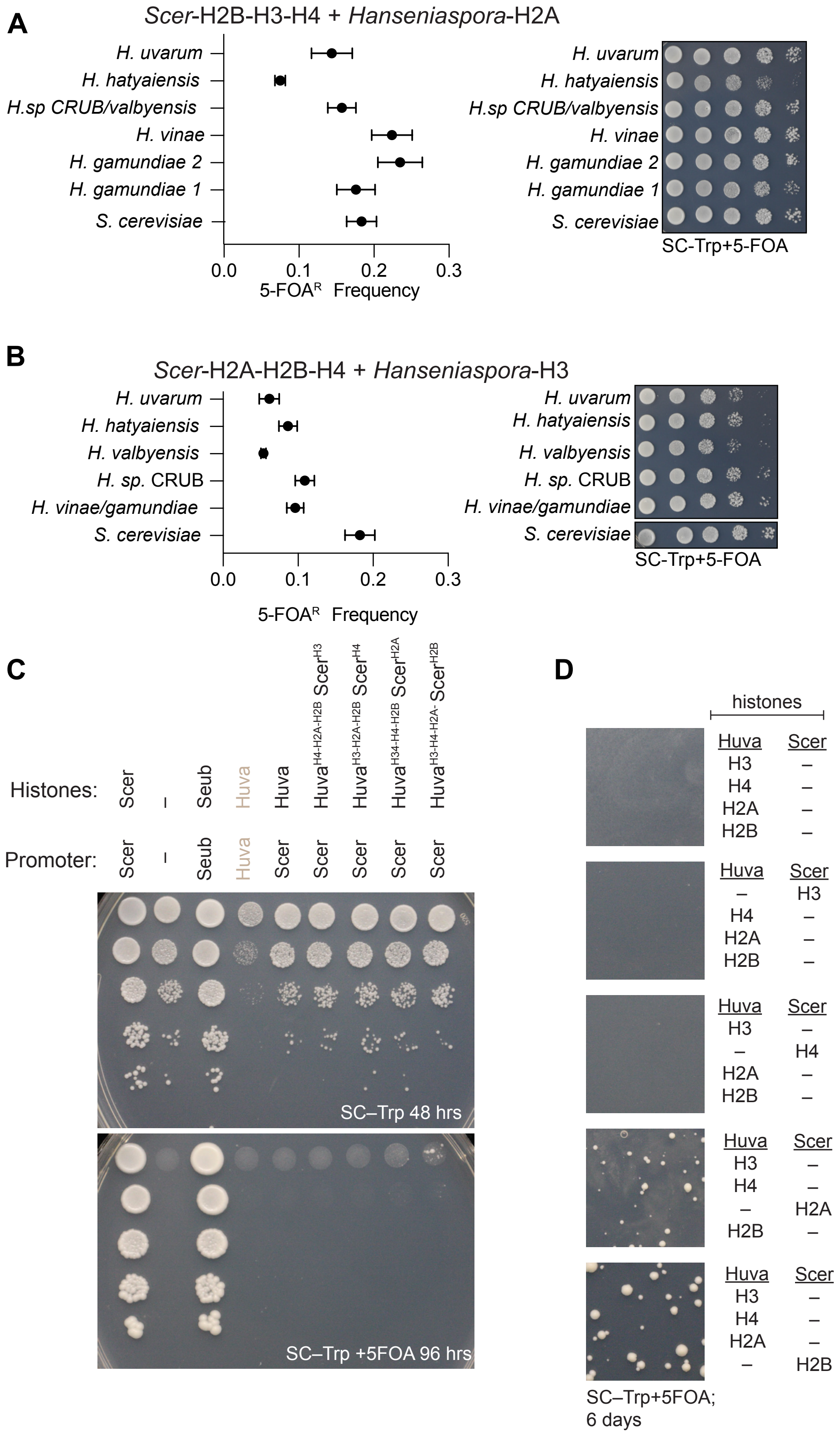

Supplementary Figure 5. Diversity of core histone cis-regulatory mechanisms in basal lineages

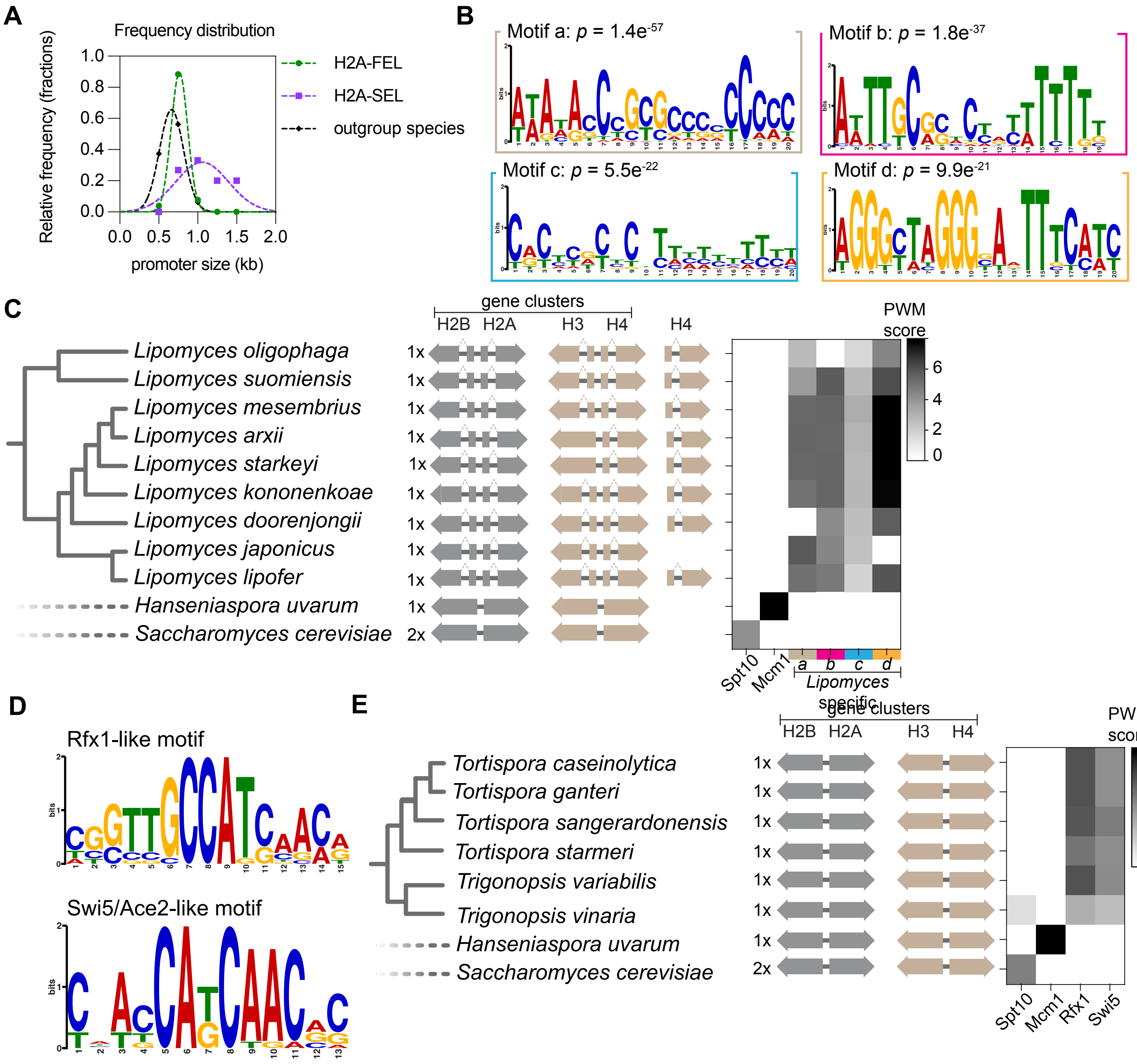

Supplementary Figure 6. Mcm1 targets genes are conserved in *H. uvarum*

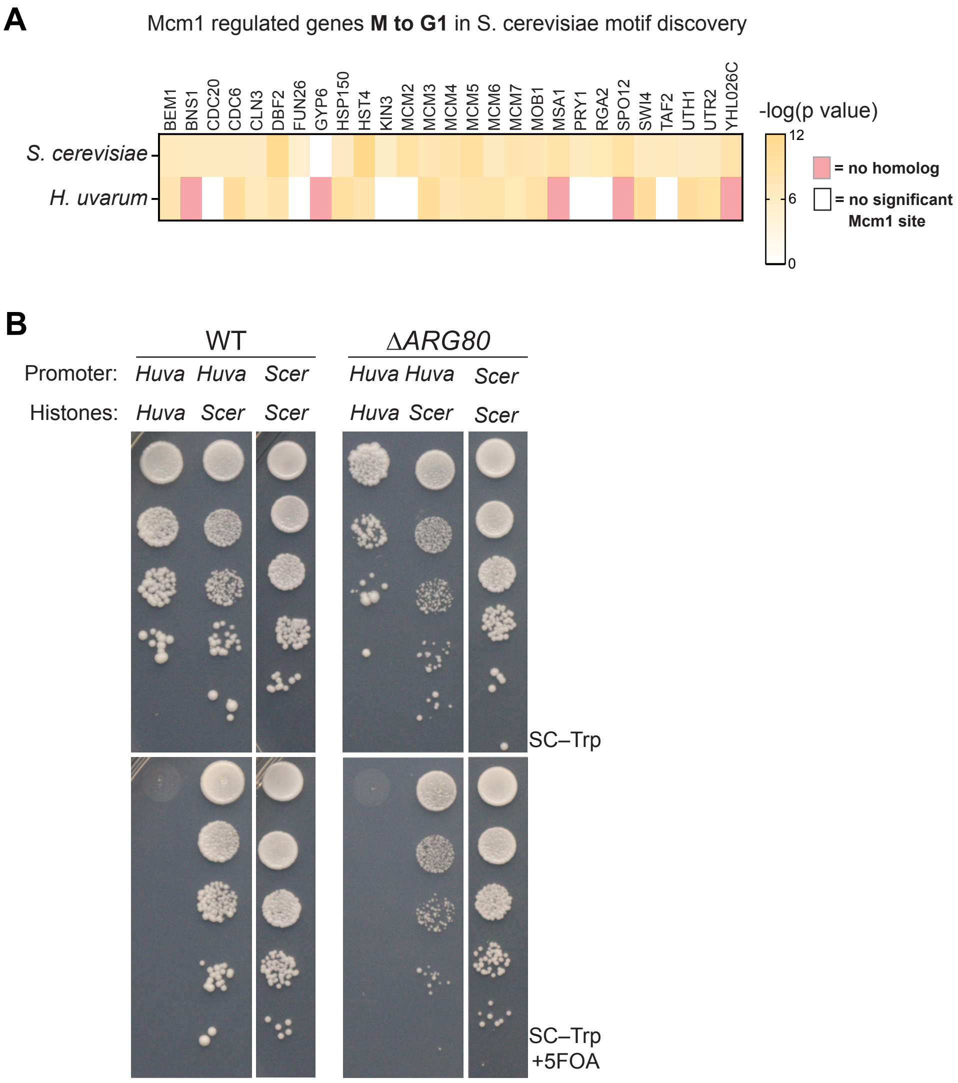

Supplementary Figure 7. Editing the Mcm1 MADS-box domain

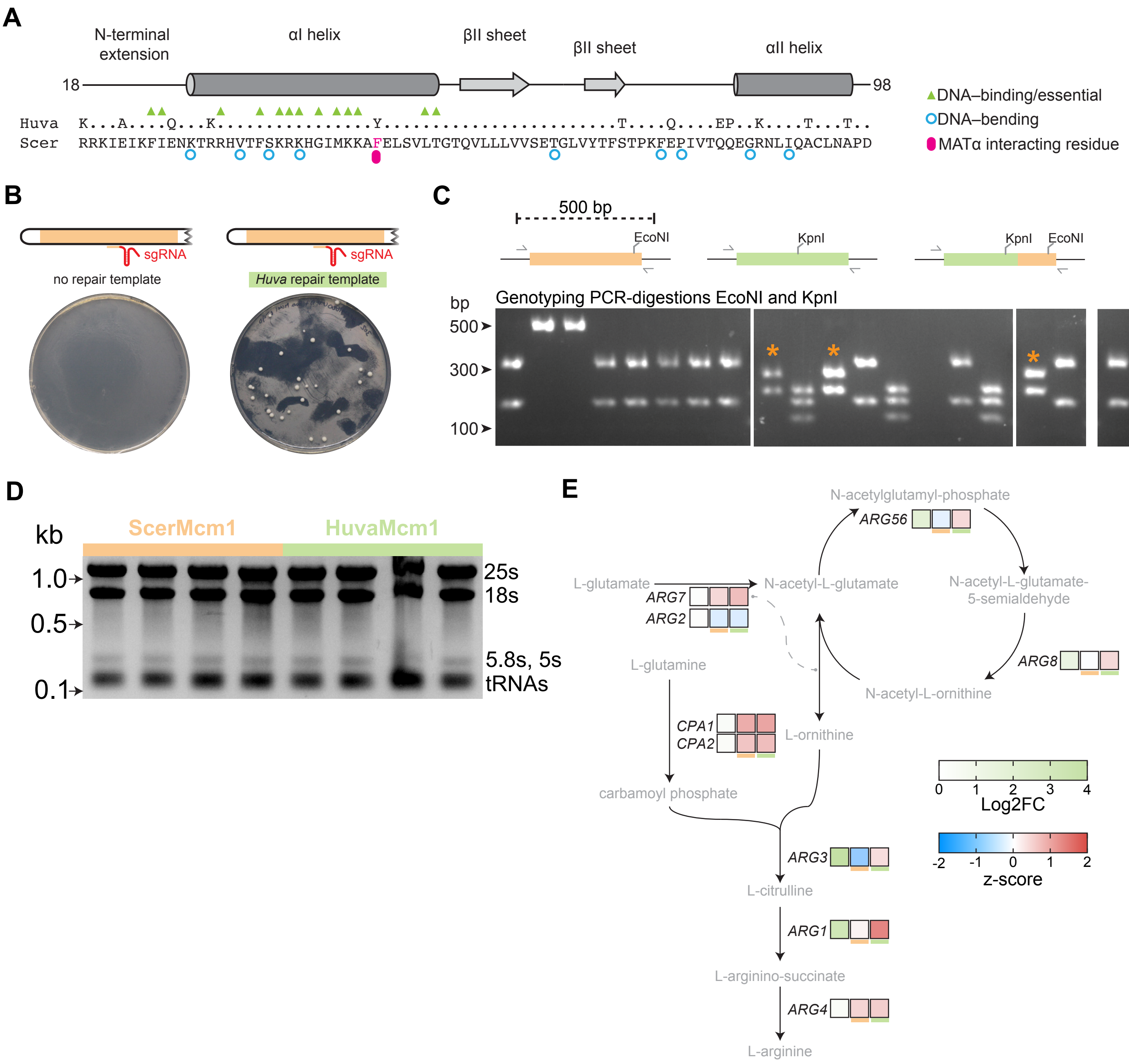

Supplementary Figure 8. Cell cycle length in *H. uvarum*

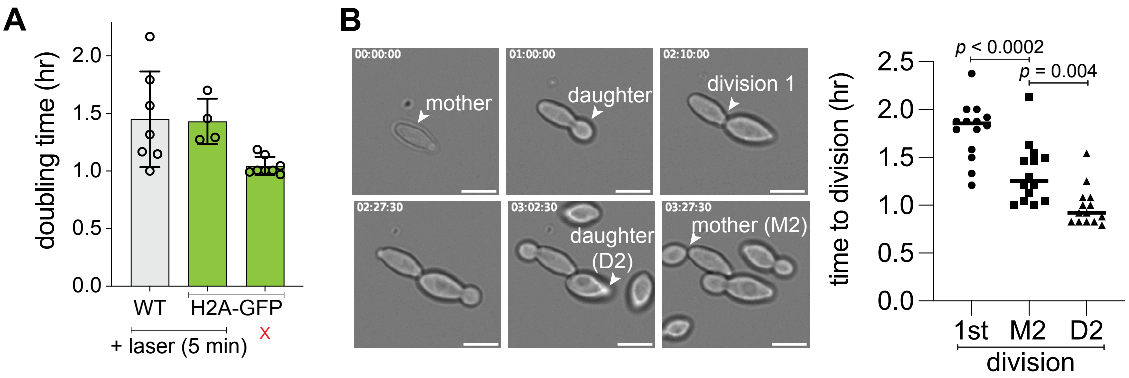
